# SCD inhibition preferentially eradicates AML displaying high de novo fatty acid desaturation and synergizes with chemotherapy

**DOI:** 10.1101/2023.08.02.551656

**Authors:** Vilma Dembitz, Hannah Lawson, Richard Burt, Céline Philippe, Sophie C. James, Samantha Atkinson, Jozef Durko, Lydia M. Wang, Joana Campos, Aoife M. S. Magee, Keith Woodley, Michael Austin, Ana Rio-Machin, Pedro Casado-Izquierdo, Findlay Bewicke-Copley, Giovanny Rodriguez Blanco, Diego Pereira Martins, Lieve Oudejans, Emeline Boet, Alex von Kriegsheim, Juerg Schwaller, Andrew J. Finch, Bela Patel, Jean-Emmanuel Sarry, Jerome Tamburini, Jan Jacob Schuringa, Lori Hazlehurst, John A. Copland, Mariia Yuneva, Barrie Peck, Pedro Cutillas, Jude Fitzgibbon, Kevin Rouault-Pierre, Kamil Kranc, Paolo Gallipoli

## Abstract

Identification of specific and therapeutically actionable vulnerabilities in acute myeloid leukaemia (AML) is needed to improve patients’ outcome. These features should be ideally present in many patients independently of mutational background. Here we identify *de novo* fatty acid (FA) desaturation, specifically stearoyl-CoA desaturase (SCD) inhibition, as a therapeutic vulnerability across multiple AML models *in vitro* and *in vivo*. We use the novel clinical grade SCD inhibitor SSI-4 to show that SCD inhibition induces AML cell death *via* pleiotropic effects, and sensitivity is based on their dependency on FA desaturation regardless of mutational profile. SSI-4 efficacy is enhanced by driving FA biosynthesis *in vitro* while stroma confers protective effects that extend to *in vivo* models. SCD inhibition increases DNA damage and its combination with standard DNA damage-inducing chemotherapy prolongs survival in aggressive murine AML models. Our work supports developing FA desaturase inhibitors in AML while stressing the importance of identifying predictive biomarkers of response and biologically validated combination therapies to realize their therapeutic potential.

**One Sentence Summary:** SCD inhibition is toxic to AML cells with high rates of fatty acid desaturation and in combination with chemotherapy prolongs survival in murine AML models.

## INTRODUCTION

AML is a highly aggressive malignant clonal disease of hematopoietic origin. Despite the approval of several novel therapies in the past decade, AML prognosis remains poor with long-term survival rates of about 30%(*1*). Development of novel therapeutic approaches for AML is particularly challenging due to high genetic and cellular heterogeneity(*2*). Therefore, identification of specific AML biological features beyond genetic mutations is needed for development of targeted therapies to improve patient outcomes. Rewired metabolism is one such feature(*3*), however discerning specific metabolic dependencies of malignant cells is crucial to avoid generalized toxicity that often compromises clinical use of metabolic inhibitors(*4*).

Fatty acid (FA) metabolism has emerged as a cancer-specific vulnerability in multiple solid cancers(*5–8*), but its role in hematological malignancies, and specifically AML, is less characterized. In AML most preclinical evidence has focused on the role of fatty acid oxidation (FAO)(*9–11*). However targeting FAO is associated with the risk of cardiac toxicity(*12*) and the best characterised FAO inhibitor, etomoxir, proved to be systemically toxic, halting its clinical development(*13*). Comparatively, targeting fatty acid synthesis (FAS), particularly stearoyl-CoA desaturase 1 (SCD1, hereafter SCD), the enzyme converting saturated fatty acids (SFA) palmitate and stearate into monounsaturated fatty acids (MUFA) palmitoleate and oleate(*14*), appears to be more tolerable based on preclinical studies(*15*). SCD plays a role in metabolic adaptation of acute lymphoblastic leukaemia to the central nervous system microenvironment(*16*) and resistance of AML stem cells to NAMPT inhibitors(*17*). Additionally, transcription factor C/AAT-enhancer binding protein α (C/EBPα) regulates lipid biosynthesis in *FLT3*-mutant AML cells by acting on the fatty acid synthase (FASN)-SCD axis, which increases their ability to withstand oxidative stress and leads to greater sensitivity to FLT3 inhibitors upon SCD inhibition(*18*). However, the significance of SCD levels in AML prognosis and response to therapy, its functional role, and potential as a therapeutic target, including identification of biological determinants of sensitivity to its inhibition, remain unknown.

Here we address these questions utilising SSI-4, a novel clinical grade SCD inhibitor with a favorable general toxicity profile(*19*). We show that SCD expression is prognostic in AML, and its inhibition compromises viability in AML cell lines and primary samples with higher rates of MUFA synthesis, without inducing hematological toxicity in mouse models. SCD activity regulates sensitivity to anthracycline-based treatments and SCD inhibition combined with standard AML chemotherapy prolongs survival in murine models of AML.

## RESULTS

### SCD levels are prognostic in AML and its inhibition induces cell death in a subset of AML cell lines

Analysis of multiple independent gene expression profiles of newly diagnosed AML samples shows that higher levels of *SCD* expression correlate with significantly decreased survival even after correcting for age, gender and European Leukemia Net (ELN) prognostic group (Fig.1A-B). This finding was confirmed when analyzing a local cohort of patients with adverse risk AML (Suppl.Fig.1A) and, after our initial report(*20*), independently validated by other research groups(*21*). High *SCD* expression correlates with several genes involved in FAS and desaturation such as fatty acid synthase (*FASN)*, fatty acid desaturase 1 and 2 (*FADS1* and *FADS2)* and adverse prognostic features such as *TP53* mutant signatures and the leukemic stem cell signature – LSC17(*22*) (Fig.1C, Suppl.Fig.1B). A biosynthesis of unsaturated fatty acids (FA) gene signature is enriched in matched post-chemotherapy relapse versus diagnosis in human AML samples (Fig.1D, Suppl.Fig.1C), together with a TCGA-generated SCD signature (Fig.1E). Interestingly, SCD signature was inversely correlated with tumor burden in cytarabine (Ara-C)-treated PDX models(*11*) and was significantly upregulated at nadir after chemotherapy (minimal residual disease, MRD) (Fig.1F). Overall these data suggest that synthesis of unsaturated FA is more active at points of leukemia resistance or progression raising the possibly that SCD is linked with mechanisms regulating sensitivity to chemotherapy. To test SCD as a therapeutic target, we used the clinical grade SCD inhibitor SSI-4 against a panel of human AML cell lines. SSI-4 induced cell death in several cell lines (K562, MOLM-13, MV-4-11), while others (OCI-AML3, THP-1, HL-60, Kasumi-1, TF-1) were resistant (Fig.1G). SSI-4 anti-leukemic effects were a result of cell death induction, as no effects on cell cycle progression were observed and sensitivity to SSI-4 was not associated to any mutational characteristics of the cell lines tested (Suppl.Fig.1D-E).

**Figure 1.**
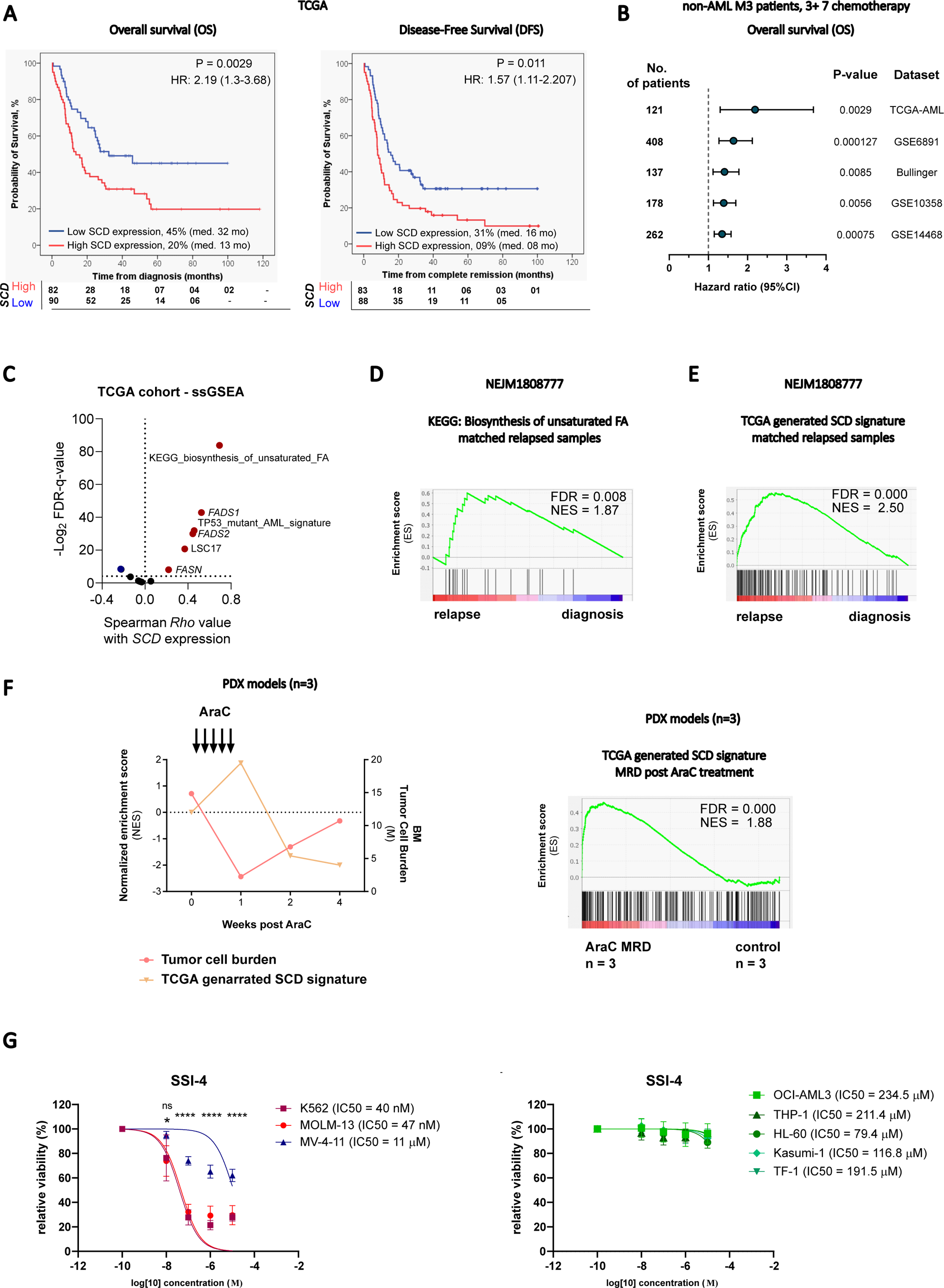
SCD expression is prognostic in AML and SCD inhibition induces cell death in a subset of AML cell lines. (A) Kaplan-Meier curves comparing overall survival and disease-free survival in TCGA AML patient cohort dichotomized after *SCD* expression. Expression level of *SCD* was considered a continuous variable and Log rank (Mantel-Cox) test was used for determining significance. (B) Forest plot of overall survival analyses considering continuous *SCD* gene expression on several patients’ datasets. Multivariate analysis corrected for confounding variables like age, gender and ELN prognostic group in all datasets. (C) Single sample gene set enrichment analysis (ssGSEA) on TCGA cohort in dependency to *SCD* expression. (D) Gene set enrichment analysis (GSEA) for KEGG pathway Biosynthesis of unsaturated fatty acids and (E) TCGA-generated SCD signature in paired diagnosis-relapse primary AML samples (NEJM1808777 dataset). (F) Tumor burden and SCD signature expression in Ara-C-treated PDX models (GSE97631) and GSEA for SCD signature at MRD stage (E) A panel of eight AML cell lines was treated with SSI-4 (0.01 - 10 µM) or corresponding vehicle for 72h. Cells with less than 10% decrease in viability were designated resistant. Results are presented as non-linear regression of normalized response and data points are mean ± SEM. * p < 0.05, ** p < 0.01, *** p < 0.001, **** p < 0.0001.

**Table 1.**
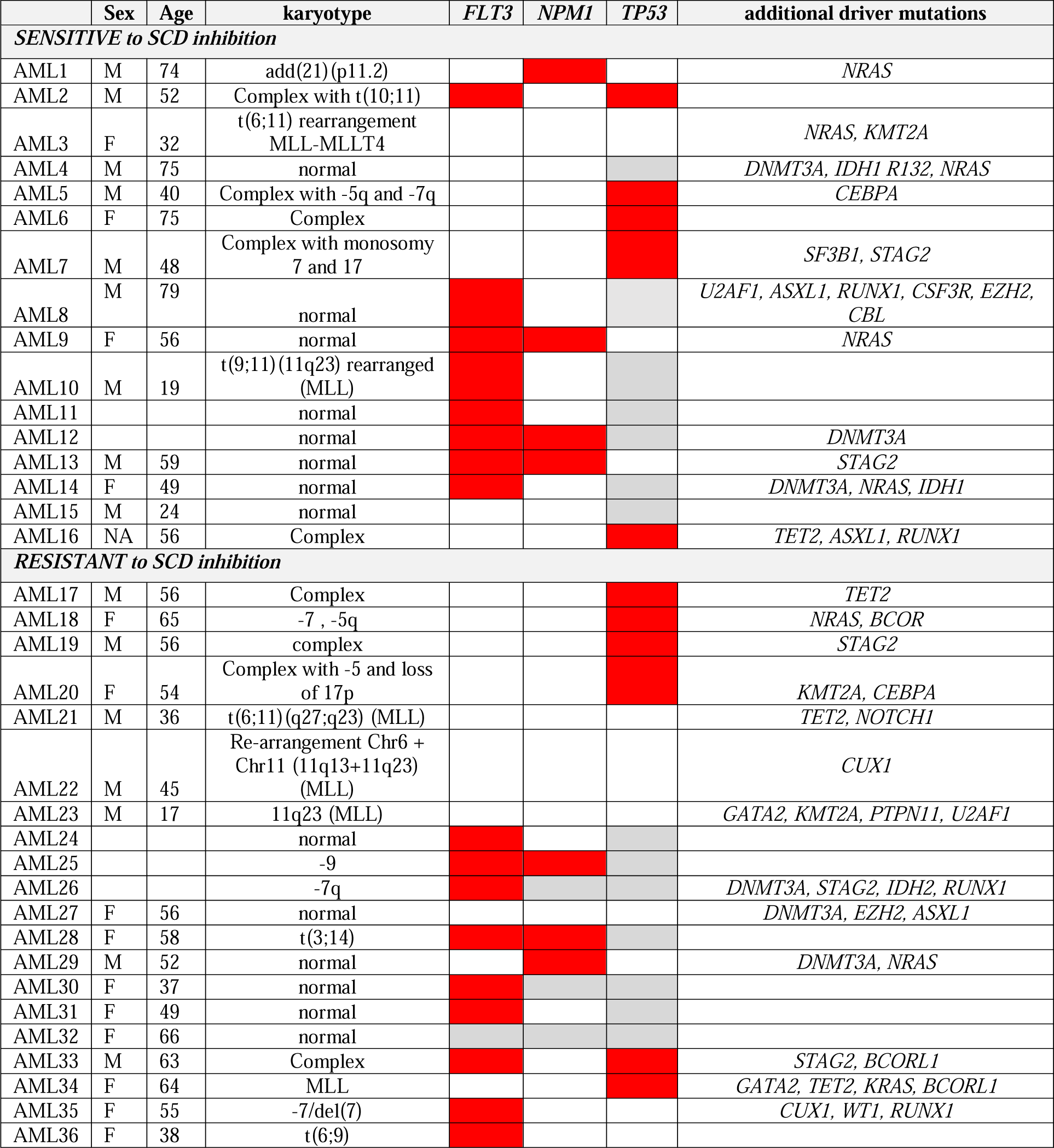
Molecular characteristics of patients samples at diagnosis (Barts Cancer Institute Tissue bank) – mutations marked as red, NA as grey, WT as white.

We validated these data using another SCD inhibitor A939572 (Suppl.Fig.2A). Moreover, SCD genetic depletion rapidly impaired growth in SSI-4-sensitive cells (Suppl.Fig.2B) but did not decrease cell viability in standard culture conditions, possibly due to compensatory upregulation of other fatty acid desaturases(*23*). However, when cells were grown in hypoxic conditions, which are known to increase FAS and dependency on SCD(*24*), SCD depletion induced cell death specifically in SSI-4 sensitive cells (Suppl.Fig.2C-D).

**Figure 2.**
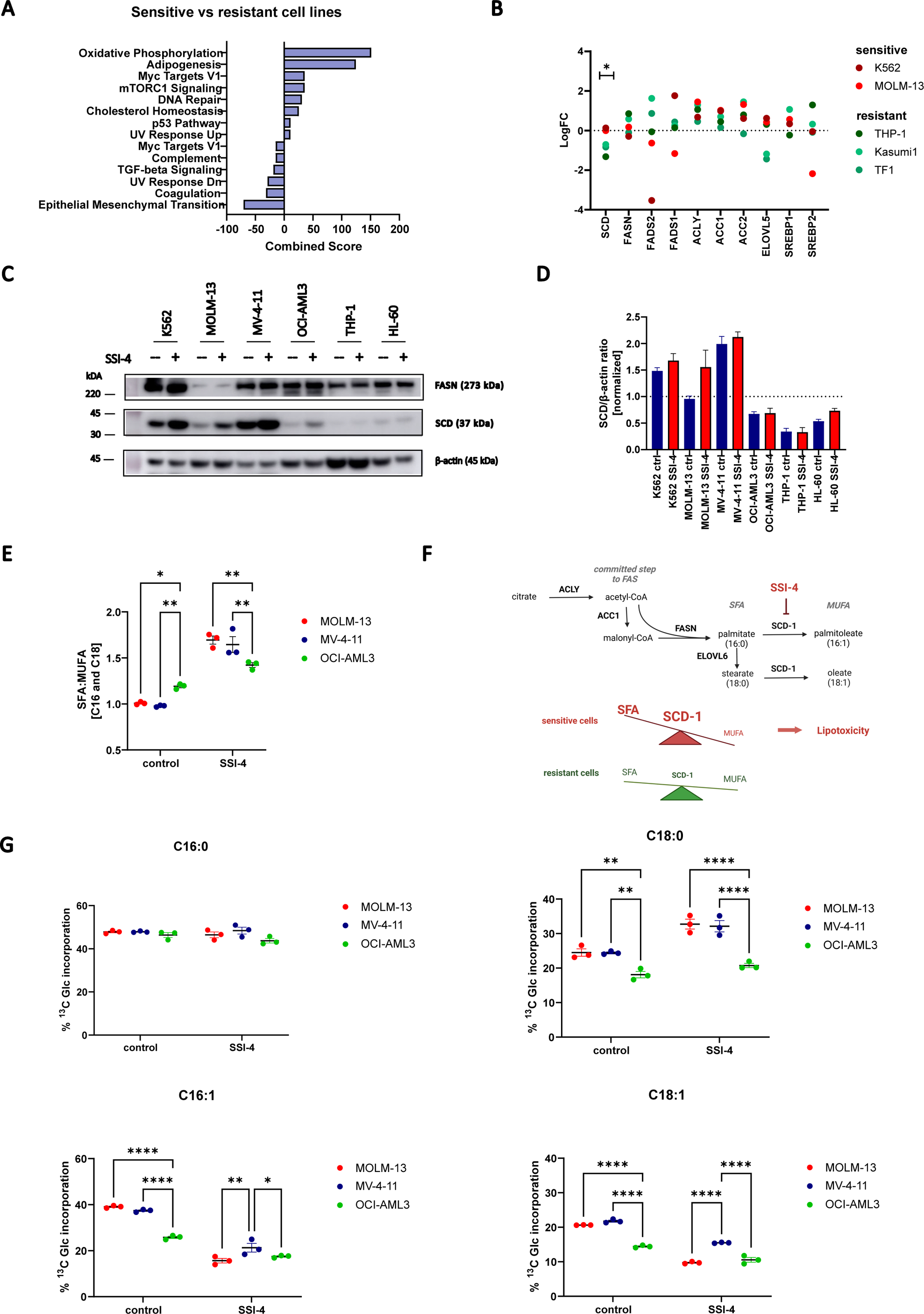
SSI-4 sensitive cells display higher levels of SCD expression and enzymatic activity. (A) Significantly enriched MSigDB signatures in sensitive vs resistant cell lines from Cancer Cell Line Encyclopedia proteomics dataset ranked by combined score from *Enrichr* enrichment analysis. Significantly upregulated signatures in sensitive cells are presented on the right hand-side and down-regulated signatures on the left hand-side of the graph. (B) Normalized expression of fatty acid synthesis related proteins in AML cell lines tested. (C) Representative western blot of sensitive (K562, MOLM-13, MV-4-11) and resistant (OCI-AML3, THP-1, HL-60) cell lines treated with SSI-4 (1 µM) or vehicle control for 24h. (D) Densitometric analysis shows SCD expression normalized to ß-actin as loading control. (E) MOLM-13, MV-4-11 and OCI-AML3 were treated with SSI-4 (1 µM) or vehicle control for 24h. Graph represents ratio of C16 and C18 saturated (SFA) and monounsaturated fatty acids (MUFA). (F) Schematic representation of *de novo* fatty acid synthesis pathway and SFA/MUFA imbalance upon SCD inhibition in sensitive and resistant cells. ACLY - ATP-citrate lyase, ACC1 – acetyl-CoA carboxylase, FASN – fatty acid synthase, ELOVL6 - ELOVL fatty acid elongase 6, SCD-1 – stearoyl-Co desaturase. (G) MOLM-13, MV-4-11 and OCI-AML3 were grown in medium supplemented with ^13^C-Glucose (2 g/L) and treated with SSI-4 (1 µM) or vehicle control for 24h. Graphs represents percentage of ^13^C-glucose incorporation in palmitate (C16:0), stearate (C18:0), palmitoleate (C16:1) and oleate (C18:1). Data are mean ± SEM. * p < 0.05, ** p < 0.01, *** p < 0.001, **** p < 0.0001.

### SCD activity dictates sensitivity to SCD inhibition by preventing SFA accumulation and lipotoxicity

Since genetic features did not correlate with sensitivity to SCD inhibition, we analyzed publicly available proteomic data(*25*) for SSI-4 sensitive and resistant cell lines. These showed an enrichment in adipogenesis signature and higher SCD protein expression in sensitive cells (Fig.2A-B). Western blot analysis confirmed these findings, and although we cannot directly correlate protein expression with enzymatic activity, this was suggestive of a greater dependency on fatty acid desaturation in sensitive cells (Fig.2C-D). Indeed fatty acid quantitation showed that the SFA/MUFA ratio is significantly higher in resistant cell lines when compared to sensitive ones. Moreover, the expected increase in ratio upon SSI-4 treatment is much lower in the resistant OCI-AML3 compared to the sensitive cells, indicating lower dependency on SCD activity in resistant cells (Fig.2E-F). Conversely sensitivity to SCD inhibition did not correlate with uptake of external lipids, or expression of lipid transporters CD36 and LDLR (Suppl.Fig.2E), both previously identified as independent prognostic factors in AML(*26, 27*). Gas chromatography-mass spectrometry (GC/MS) experiments tracing uniformly-labelled ^13^Carbon (U-^13^C_6_) glucose incorporation into FA demonstrated that in sensitive cells MUFA biosynthesis (16:1, C18:1) was higher and more responsive to SCD inhibition. Interestingly, treatment with SSI-4 also led to increased production of SFA stearate (C18:0) in sensitive cells. Together these data suggest that SSI-4 sensitive cells have a more active *de novo* fatty acid biosynthesis and desaturation. Moreover upon SCD inhibition FAS is further increased but becomes uncoupled from desaturation thus causing an imbalance between SFA and MUFA levels to a degree able to trigger lipotoxicity(*28*) and cell death (Fig.2F-G).

### MUFA and SFA levels modulate sensitivity to SCD inhibition in AML cells by regulating FAS

The above data suggest that, in cells sensitive to SCD inhibition, MUFA production is a sensing point for FAS pathway activity. To test this hypothesis, we grew cells in the presence of oleate and noted decreased production of both SFA and MUFA (Fig.3A and Suppl.Fig.3A-B) which also correlated with a complete rescue of SSI-4-mediated decrease in viability, supporting the on-target efficacy of SSI-4 (Fig.3B). Conversely, addition of palmitate in non-toxic concentration increased FAS and oleate production mostly in resistant, but not sensitive cells (Fig.3C and Suppl.Fig.3A-B). This suggests that baseline MUFA production in resistant cells is below its potential maximum and can be upregulated following palmitate supplementation. Although exogenous palmitate can still be detoxified by desaturation both in sensitive and resistant cells, as shown by total levels of labelled and unlabeled MUFA (Suppl.Fig.3B), in palmitate rich conditions SCD inhibition results in a large increase in the SFA/MUFA ratio. This is true even in resistant cells that do not display such strong imbalance upon treatment with SSI-4 alone (Fig.2E, Suppl.Fig.3A-C) and causes increased sensitivity to SSI-4 in both sensitive and resistant cells in the presence of palmitate (Fig.3D).

**Figure 3.**
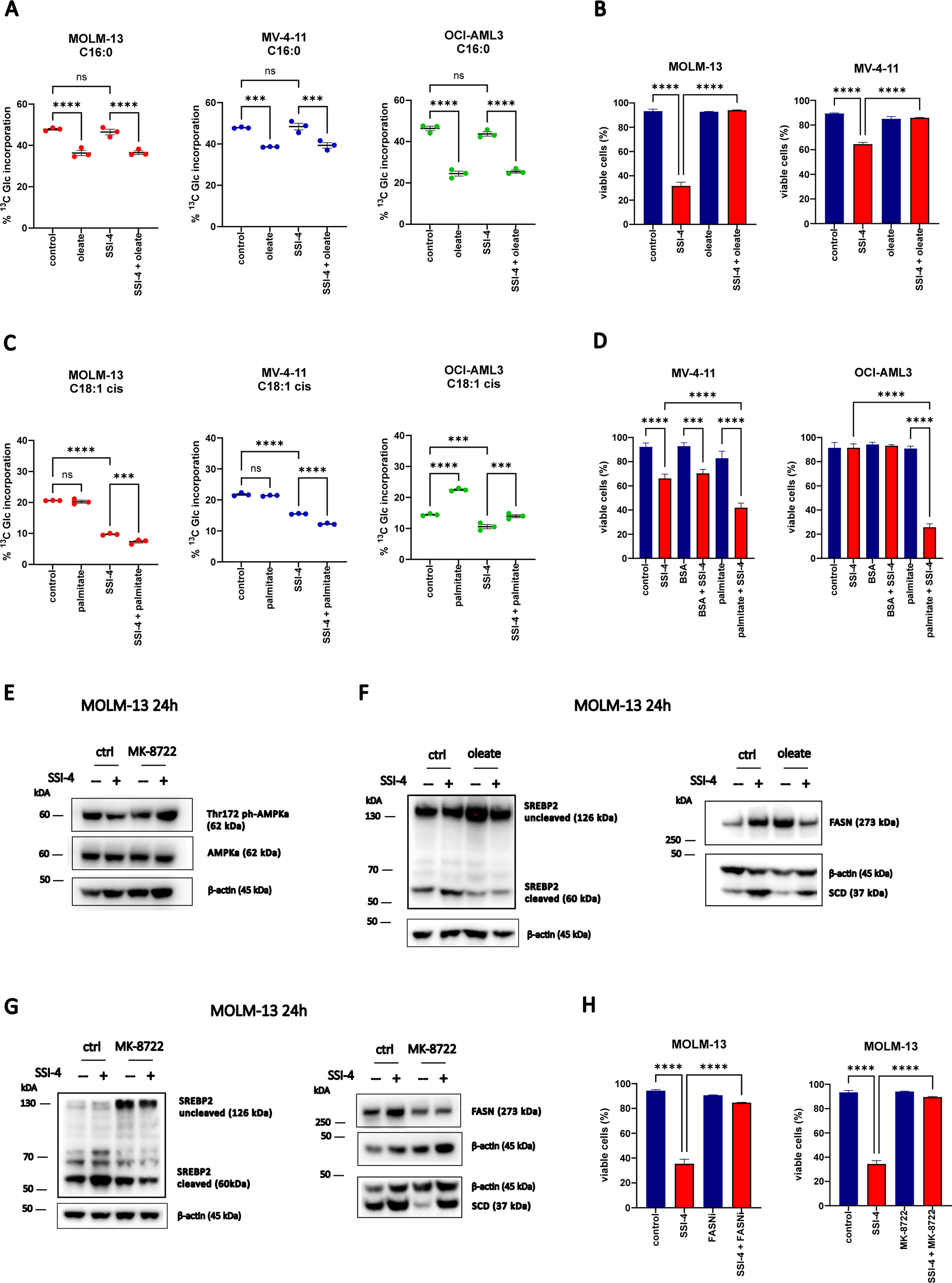
SFA/MUFA ratio and lipotoxicity in response to SCD inhibition depend on the activity of FAS pathway. (A, C) MOLM-13, MV-4-11 and OCI-AML3 were labelled with ^13^C-Glucose (2 g/L) and treated for 24h with SSI-4 (1 µM) or vehicle control with or without the addition of oleate (100 µM) or palmitate (100 µM). Graphs represents percentage of ^13^C-glucose incorporation in palmitate (C16:0) and oleate (C18:1) (B, D) MOLM-13, MV-4-11 and OCI-AML cells were treated for 72h with SSI-4 (1 µM) with or without addition of oleate (100 µM) or palmitate (100 µM). (E-G) Representative western blots of MOLM-13 cells treated for 24h with SSI-4 (1 µM) or vehicle control with or without addition of oleate (100 µM) or direct AMPK activator MK-8722 (10 µM). (H) MOLM-13 cells were treated for 72h with SSI-4 (1 µM) with or without addition of FASN inhibitor Fasnall (20 µM) or MK-8722 (10 µM). Viable cells were determined as Annexin-V^-^/PI^-^. Data are mean ± SEM.* p < 0.05, ** p < 0.01, *** p < 0.001, **** p < 0.0001.

These data demonstrate that changes in MUFA/SFA levels affect sensitivity to SCD inhibition via modulation of FAS. To clarify the mechanistic underpinning of this observation we analyzed changes in regulatory pathways of FAS in sensitive cells. Consistent with the observed increase in the FAS rate, SCD inhibition reduced activation of AMPK (Fig.3E), thereby relieving its inhibitory role on cleavage and consequent activation of SREBP2 (Fig.3F), a key transcription factor modulating FAS enzymes expression(*29, 30*). As expected following SREBP2 activation, we observed increased levels of FASN, SCD (Fig.2C and 3F) and total acetyl-Coa carboxylase (ACC) which were reversed by the addition of oleate, thus confirming that MUFA levels relieve SCD inhibition toxicity through downregulation of FAS (Fig.3F, Suppl.Fig.4A). Conversely, the AMPK activator MK-8722 decreased both SREBP2 cleavage and expression of SCD and FASN (Fig.3G). Although SREBP2 has mostly been described as a regulator of cholesterol synthesis, while FAS is generally under regulation of SREBP1(*30*), in our system we observed more consistent effects on SREBP2 following SCD inhibition and MUFA addition. Conversely, we did not see a change in SREBP1 cleavage in response to SSI-4, with or without the addition of oleate, even though AMPK activation with MK-8722 decreased SREBP1 cleavage as expected (Suppl.Fig.4B). These data suggest that FAS in AML cell lines is prominently regulated by SREBP2, as shown in other models(*31*).

**Figure 4.**
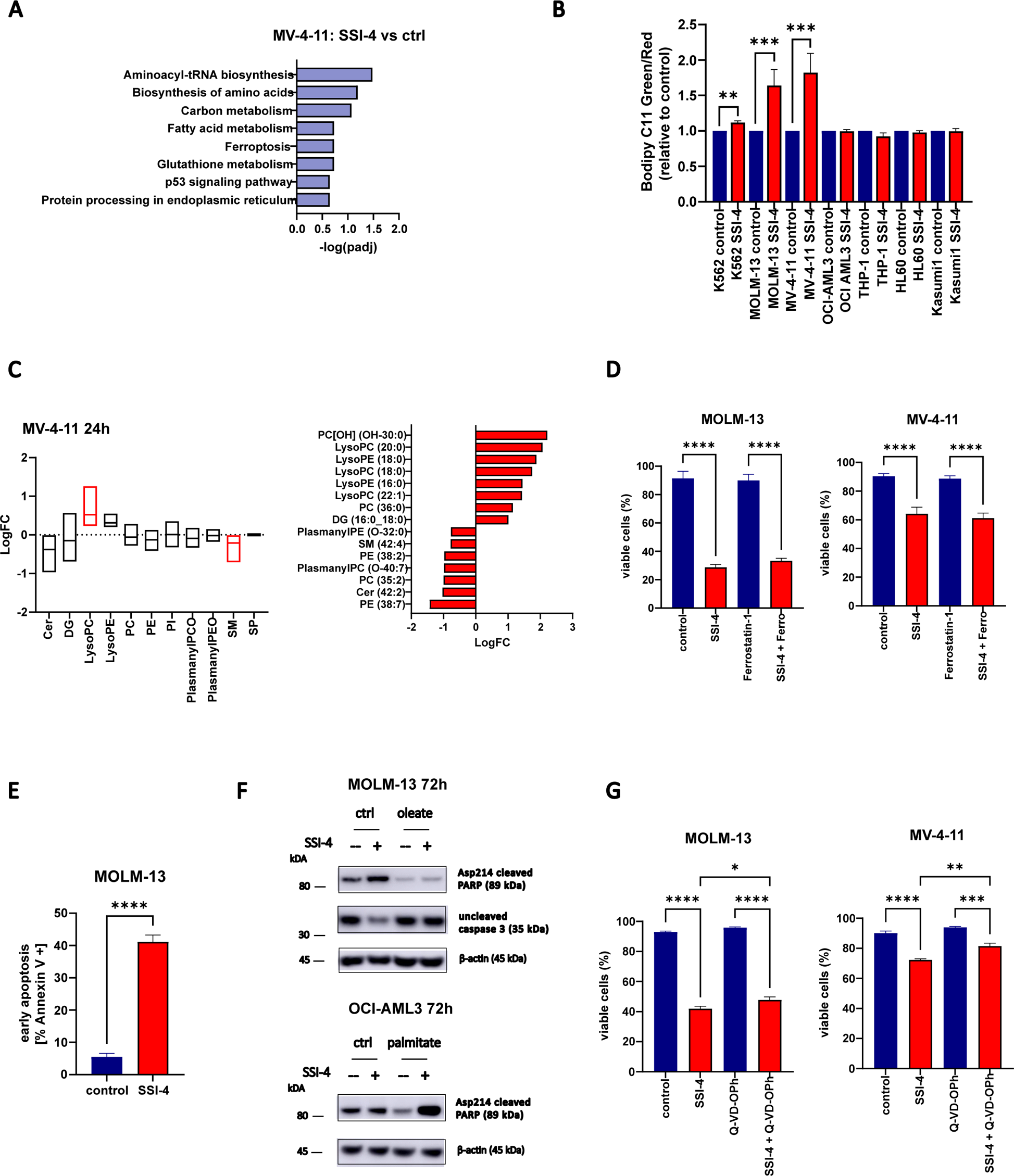
SSI-4 treatment induces both an increase in lipid peroxidation and activation of apoptotic machinery. (A) Significantly enriched KEGG pathway signature in MV-4-11 cells treated with SSI-4 (1 µM) or vehicle control for 24h. (B) Sensitive K562, MOLM-13, MV-4-11 and resistant OCI-AML3, THP-1, HL-60 and Kasumi-1 cell were treated for 24h with SSI-4 (1 µM) or vehicle control. Lipid peroxidation was measured using Bodipy C11. (C) Lipidomics analysis on MV-4-11 cells treated for 24h with SSI-4 (1 µM) or vehicle control. Upper graph represents enrichment analysis per lipid groups of treated cells vs. control (Q1-Q3 with line at median value) with significant lipid groups marked in red. Lower graph represents significant differentially expressed individual lipids with upregulated lipids presented on the right hand-side and down-regulated lipids on the left hand-side of the graph. Red bars: padj < 0.05. (D) MV-4-11 and MOLM-13 cells were treated for 72h with SSI-4 (1 µM) or vehicle control with or without addition of Ferrostatin-1 (5 µM). (E) MOLM-13 cells in early apoptosis (Annexin-V^+^/PI^-^) after 72h treatment with SSI-4 (1 µM). (F) Representative western blot of MOLM-13 and OCI-AML3 cells treated for 72h with SSI-4 (1 µM) or vehicle control with or without addition of oleate (100 µM) or palmitate (100 µM). (G) MOLM-13 and MV-4-11 cells were treated for 72h with SSI-4 (1 µM) or vehicle control with or without addition of Q-VD-OPh (50 µM). Viable cells were determined as Annexin-V^-^/PI^-^. Data are mean ± SEM. * p < 0.05, ** p < 0.01, *** p < 0.001, **** p < 0.0001.

Decreasing FAS by inhibition of either FASN or ACC or via AMPK activation abolished SSI-4 mediated toxicity (Fig.3H, Suppl.Fig.4B-C). Interestingly, analysis of the Depmap dataset shows that SCD dependency inversely correlates with the expression levels of both *FASN* and *ACACA* (ACC) across all cancer cells lines and particularly AML ones (Suppl.Fig. 4D). Overall these data confirm that modulation of FAS impacts sensitivity to SCD inhibition.

### Cell death in response to SSI-4 is mediated by lipid oxidative stress, integrated stress response and activation of apoptotic machinery

Transcriptomic analysis of SSI-4 treated cells confirmed the regulatory role of oleate levels on the rate of FAS but also identified oxidative stress associated pathways (ferroptosis, glutathione metabolism) and integrated stress/endoplasmic reticulum (ER) stress response as potential downstream mechanisms leading to cell death (Fig.4A and Suppl.Fig.5A).

**Figure 5.**
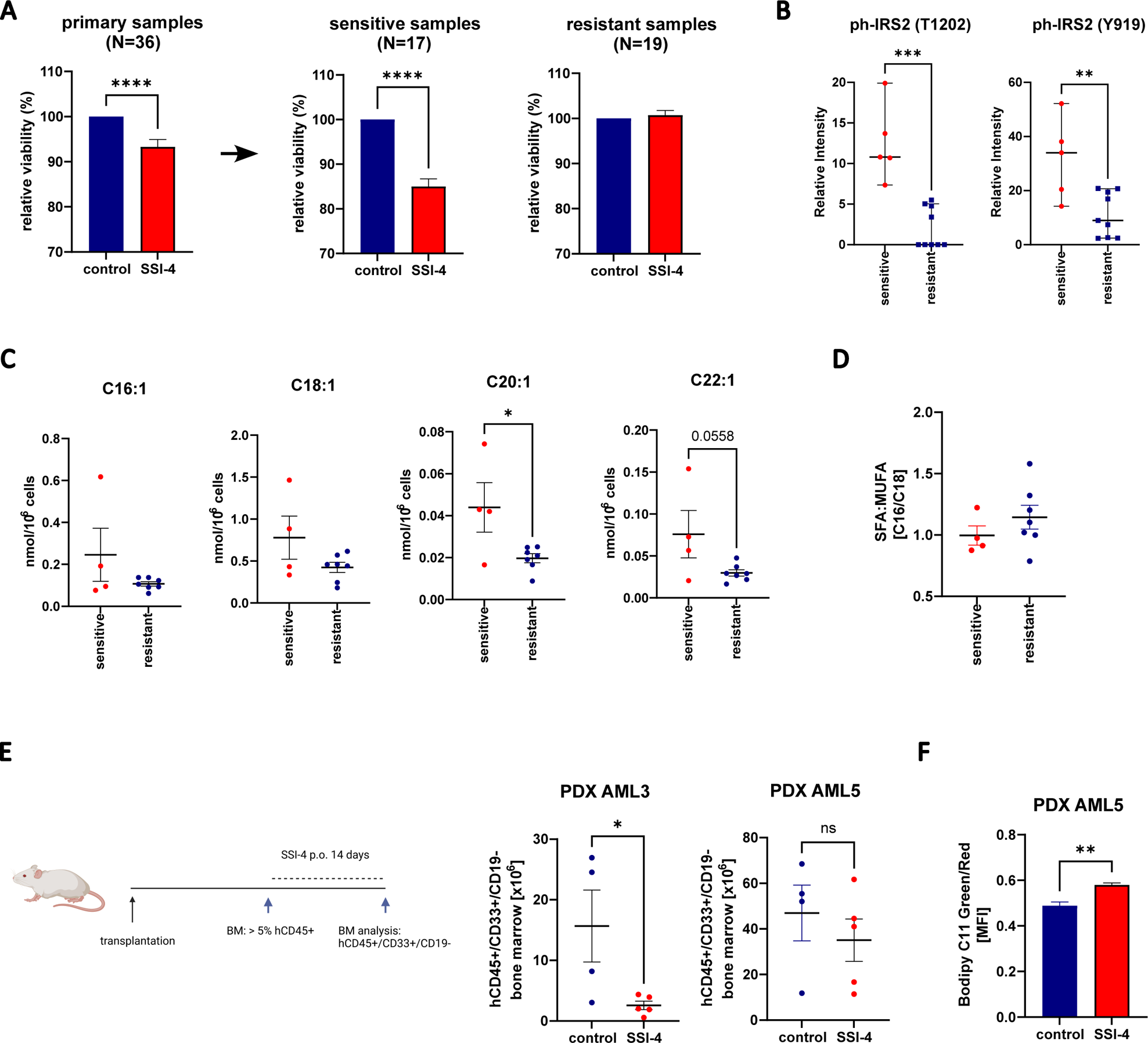
SSI-4 has antileukemic effects on a subset of primary AML patient samples both *in vitro* and *in vivo* with sensitive samples showing higher levels of MUFA. (A) AML primary samples (n=36) were depleted of T-cells and grown in co-culture with irradiated MS-5 cells for 7 days with addition of SSI-4 (1 µM). Samples with less than 5% decrease in viability were designated to resistant group. (B) Phosphorylated IRS2 in sensitive (n=5) vs resistant (n=11) samples measured using phosphoproteomic analyses. Graphs represent relative intensity of phosphopeptides. (C-D) MUFA levels and SFA/MUFA ratio in sensitive (n=4) vs resistant (n=7) samples. (E) Two sensitive samples *in vitro* were transplanted into NBSGW mice. When engraftment of human CD45^+^ cells exceeded 5% in the BM, animals were treated with 10 mg/kg of SSI-4 or corresponding vehicle orally for 14 days. Total number of human leukemic cells (hCD45^+^hCD33^+^hCD19^-^) isolated from two legs at the end of the experiment. (F) In hCD45^+^hCD33^+^ cells isolated from mice transplanted with AML5 sample lipid peroxidation was measured using Bodipy C11. Viable cells were determined as Annexin-V^-^/PI^-^. Data are mean ± SEM. * p < 0.05, ** p < 0.01, *** p < 0.001, **** p < 0.0001.

Consistent with SCD role in protection against oxidative stress and lipid peroxidation(*18*), sensitive cells displayed a specific increase in lipid peroxidation as measured by Bodipy C11 staining upon treatment with SSI-4 (Fig.4B). A similar effect was observed in cells with downregulated SCD in hypoxic condition where lipotoxicity is present (Suppl.Fig.5B). Lipidomic analysis confirmed that lysophospholipids which have lost their polyunsaturated tail, a known marker of lipid peroxidation(*32*), are the most enriched lipid class in response to SSI-4 (Fig.4C). This pattern was completely abrogated by the addition of oleate (Suppl.Fig.5C). However, despite reducing peroxidation to the same extent of oleate (Suppl.Fig.5D-E), lipid peroxidation inhibitors failed to or only partially rescued (Fig.4D and Suppl.Fig.5F-G) sensitive cells from SSI-4-mediated cell death. Reversal of lipid peroxidation is therefore not sufficient to prevent cell death induced by SCD inhibition.

SCD is essential for ER homeostasis(*33*) and SFA/MUFA imbalance is known to trigger ER stress(*34*). Based on our transcriptomic data, we interrogated the three arms of the ER response pathway (Suppl.Fig.6A). We noticed a substantial increase in targets downstream of PERK, CHOP and ATF4, a moderate increase in IRE1 and IRE1-associated targets, spliced and total XBP1, and no effects on the expression of ATF6. In accordance to that, PERK inhibitor GSK2656157 rescued SSI-4-mediated cytotoxicity, while IRE1 inhibitor 4µ8c demonstrated only milder cytoprotective effects at higher doses and ATF6 inhibitor Ceapin A7 had no effects (Suppl.Fig.6B). The PERK pathway is a known regulator of apoptosis(*35*) and SSI-4 treatment induced accumulation of apoptotic marker Annexin V and activation of apoptotic machinery in both sensitive cells and resistant ones grown in the presence of palmitate (Fig.4E-F). Still, in contrast to oleate supplementation, co-treatment with pan-caspase inhibitor Q-Vd-OPh resulted in a significant, but only partial rescue of SSI-4 mediated cell death (Fig.4F-G).

**Figure 6.**
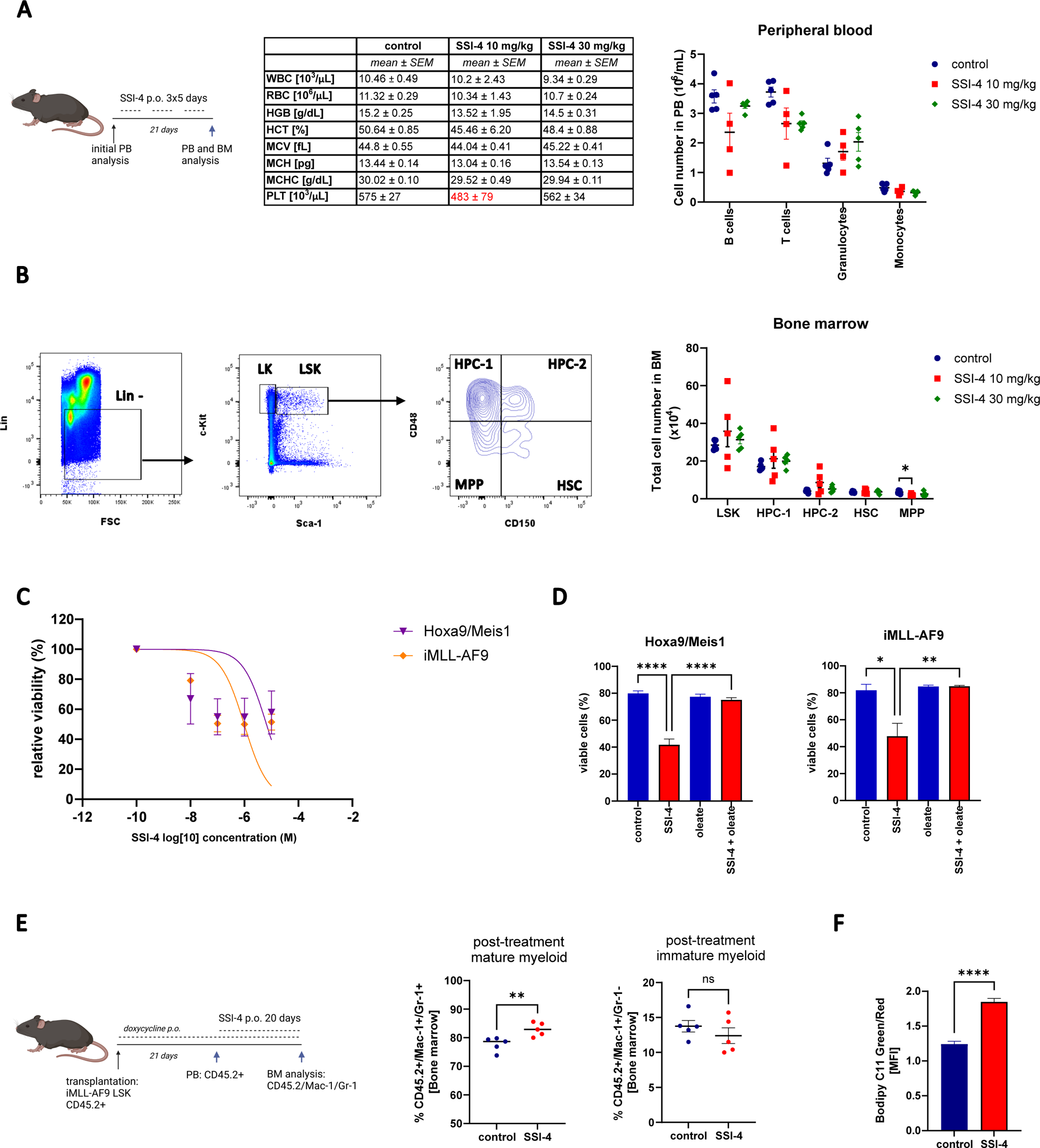
SSI-4 demonstrates no hematopoietic toxicity but, as single agent, does not prolong survival in a mouse AML model. (A) C57BL/6 mice (n=15) were treated with 10 and 30 mg/kg SSI-4 or corresponding vehicle orally for a total of 21 days with 2 days break after each 5 days of continuous treatment. PB counts of control or SSI-4 treated mice. WBC – white blood cells, RBC – red blood cells, HGB – hemoglobin concentration, HCT – hematocrit, MCV – mean cell volume, MCH – mean cell hemoglobin, MCHC – mean cell hemoglobin concentration, PLT – platelets. (B) Gating strategy and total numbers of cells in LSK (Lin^-^Sca-1^+^c-Kit^+^), HPC-1 (LSK CD48^+^CD150^-^), HPC-2 (LSK CD48^+^CD150^-^), HSC (LSK CD48^-^CD150^+^), MPP (LSK CD48^-^CD150^-^) compartments in the BM isolated from two legs of treated animals. (C) Leukemic *Hoxa9/Meis1* and *iMLL-AF9* cells were treated with SSI-4 (0.01-10 µM) or corresponding vehicle. (D) Leukemic *Hoxa9/Meis1* and *iMLL-AF9* cells were treated with SSI-4 (1 µM) or vehicle control with or without addition of oleate (100 µM) for 48h. (E) CD45.2^+^ LSK cells from *iMLL-AF9* mice were transplanted to lethally irradiated syngeneic CD45.1^+^/CD45.2^+^ recipient mice (n=10). After engraftment was confirmed in PB, animals were treated with 10 mg/kg SSI-4 or corresponding vehicle orally for 20 days. Percentage of CD45.2^+^/Mac-1^+^/Gr-1^+^ and CD45.2^+^/Mac-1^+^/Gr-1^-^ cells in the BM at the end of the experiment. (F) In CD45.2^+^ cells lipid peroxidation was measured using Bodipy C11. Viable cells were determined as Annexin-V^-^/PI^-^ or Annexin-V^-^/7-AAD^-^. Data are mean ± SEM. * p < 0.05, ** p < 0.01, *** p < 0.001, **** p < 0.0001.

Overall these data show that the lipotoxic reaction in response to SSI-4 cannot be reduced to the activation of a single effector death mechanism and that SCD inhibition act as a pleiotropic trigger which can activate several cell death modes concurrently, thus explaining the detection of both ferroptosis or apoptosis markers(*36*). Consistent with this, inhibiting any of these cell death mechanisms independently did not completely rescue the cytotoxic effects of SSI-4 supporting their functional redundance.

### Primary AML cells with higher levels of unsaturated fatty acids are sensitive to SCD inhibition

To extend the translational relevance of our findings, we tested the effects of SSI-4 on primary AML samples *in vitro*. First, we observed that co-culture of primary AML samples with MS-5 cells decreased sensitivity to SSI-4 and abolished palmitate-mediated toxicity, suggesting that the presence of stroma could influence SCD dependency (Suppl.Fig.7A). We thus treated 36 primary AML samples in stromal co-culture with SSI-4 and observed that, similar to cell lines, primary samples dichotomized into sensitive and resistant (Fig.5A), with no clear relation to specific driver mutations (Suppl. Table1). This was confirmed on an independent cohort of 11 primary samples (Suppl.Fig.7C). We did not identify a sensitivity signature using transcriptomic or proteomic approaches (data not shown), while phosphoproteomic data on a subset of AML patient samples showed significantly higher levels of phosphorylated-insulin receptor substrate 2 (IRS2) in sensitive cells (Fig.5B) which was confirmed in an independent phosphoproteomic analysis on a different patient sample subset (Suppl.Fig.7D). IRS2 is a downstream target of receptor tyrosine kinases (RTK) and has been shown to specifically regulate the insulin-like growth factor-1 (IGF-1) autocrine production and signaling in AML(*37*). The activation of RTK was further supported by increased phosphorylation of AKT2 and PLEKHG3 (Suppl.Fig.7E). Interestingly, increased activation of signaling downstream of RTK in this AML model did not correlate with faster progression through cell cycle (Suppl.Fig.7F) suggesting that sensitivity is not linked to a more proliferative phenotype, but potentially to the role of RTKs in regulation of lipogenesis and energy metabolism(*38*). Indeed, in accordance with cell line data, sensitive samples displayed higher levels of MUFAs and a lower SFA/MUFA ratio compared to resistant cells supporting their greater dependency on fatty acid desaturation (Fig.5C-D).

**Figure 7.**
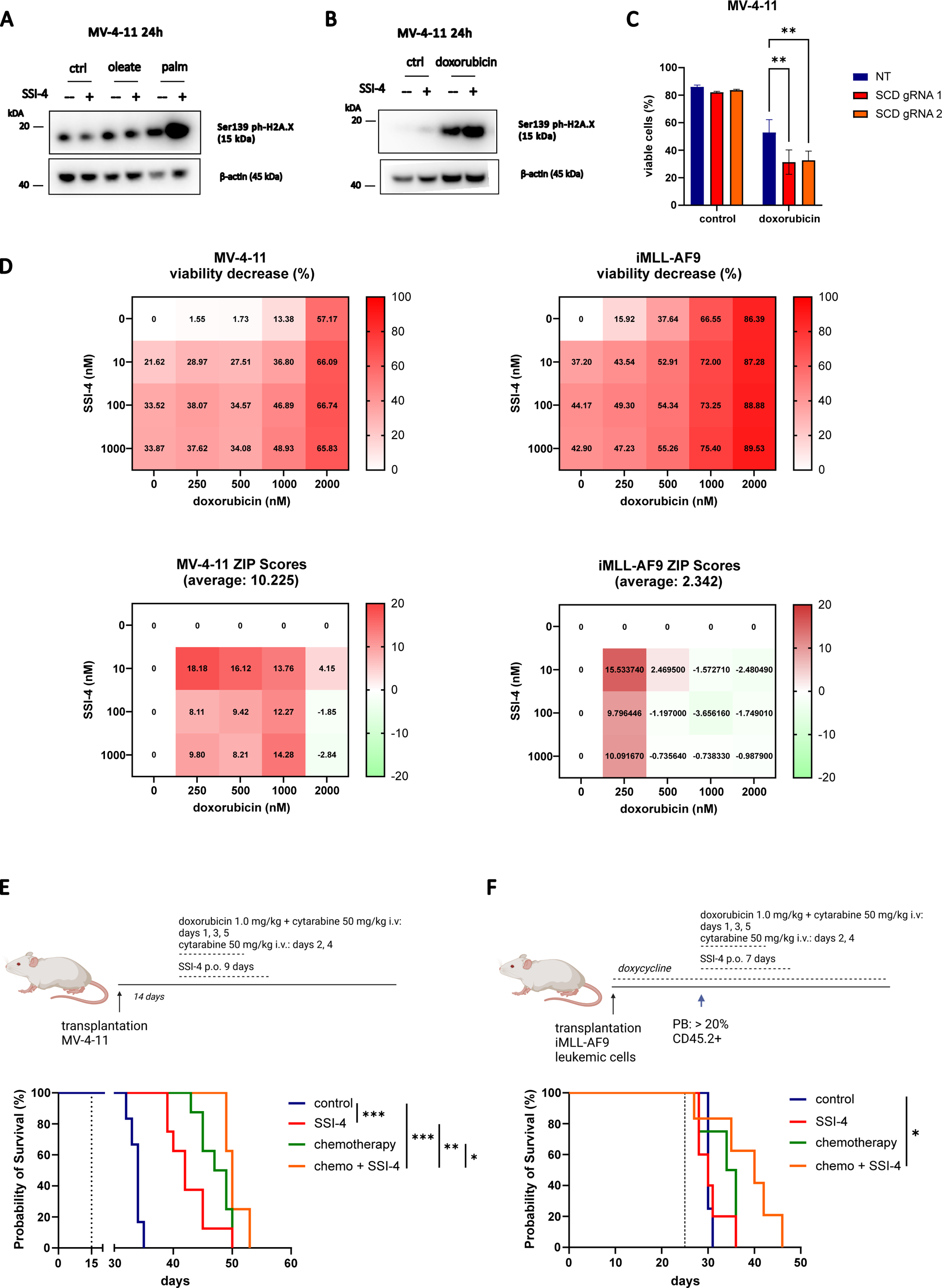
Lipotoxicity increases DNA damage and SSI-4 acts synergistically with DNA damage inducing chemotherapy both *in vitro* and *in vivo*. (A-B) Representative western blots of MV-4-11 cells treated with SSI-4 (1 µM) or vehicle control with or without addition of oleate (100 µM), palmitate (100 µM) or doxorubicin (1 µM) for 24h. (C) MV-4-11 NT gRNA, SCD gRNA 1 and SCD gRNA 2 were treated for 72h with doxorubicin (1 µM). (D) MV-4-11 and leukemic iMLL-AF9 cells were treated for 72h with growing concentrations of SSI-4 and doxorubicin. Synergy was determined by Bliss coefficient (ZIP Score > 10 indicates synergism). (E) MV-4-11 cells were transplanted into NBSGW mice (n=28). 14 days after transplant animals were treated for 9 days with 10 mg/kg SSI-4 or corresponding vehicle orally with or without conventional chemotherapy protocol (3 days 1.0 mg/kg doxorubicin i.v., 5 days 50 mg/kg cytarabine i.v.). (F) CD45.2^+^ leukemic *iMLL-AF9* cells were transplanted into CD45.1^+^ NBSGW mice (n=18). When leukemic burden in PB reached 20%, animals were treated for 7 days with 10 mg/kg SSI-4 or corresponding vehicle orally with or without conventional chemotherapy protocol. Kaplan-Meier curves represent overall survival of animals treated with SSI-4 and corresponding vehicle with our without conventional chemotherapy. Viable cells were determined as Annexin-V^-^/PI^-^ or Annexin-V^-^/Zombie^-^. Data are mean ± SEM. * p < 0.05, ** p < 0.01, *** p < 0.001, **** p < 0.0001.

Given the observed stromal protection towards SSI-4-mediated toxicity, we tested to which extent the anti-leukemic effect of SCD inhibition was maintained *in vivo* using patient derived xenografts (PDX) from two sensitive AML samples. A decrease in bone marrow (BM) leukemia burden following SSI-4 treatment was observed in both PDX model, although this reached statistical significance only in one. Still, higher levels of lipid peroxidation in the human CD45^+^ compartment were noted following SSI-4 even when leukemic burden was not significantly reduced (Fig.5E-F, Suppl.Fig.7G). This suggests that SCD inhibition primes AML cells to aberrant oxidative stress *in vivo* even when its antileukemic effects are blunted.

### In-vivo SSI-4 treatment does not affect normal hematopoiesis and induces differentiation and lipid oxidative stress in iMLL-AF9 murine AML model

Although a favorable general toxicity profile of SSI-4 was demonstrated in previous animal studies (*5, 19*), we also ascertained its potential hematopoietic toxicity. SSI-4 had no significant hematopoietic toxicity, as shown by its effects on the peripheral blood (PB) counts and hematopoietic progenitor compartments of treated animals (Fig.6A-B).

We then assessed the effects of SCD inhibition in two AML murine models expressing either the *Hoxa9/Meis1* or inducible *MLL-AF9* (*iMLL-AF9*) oncogene. Both models were sensitive *in vitro* to SSI-4 with toxicity reversed by addition of oleate (Fig.6C-D). Treatment *in vivo* of *iMLL-AF9* model with SSI-4 resulted in induction of differentiation without significant decrease in BM leukemic burden (Fig.6E). However, as seen in PDX, lipid peroxidation was increased in treated AML cells *in vivo* (Fig.6F).

### SSI-4 combination with doxorubicin based chemotherapy is synergistic and prolongs survival in murine AML models

Lipid peroxidation is the most consistent phenotype observed in response to SSI-4 across all AML models tested and is known to induce DNA damage(*39*) through production of malondialdehyde which forms DNA adducts in normal and oncogenic mammalian cells(*40*). Moreover a role of SCD inhibition in modulating DNA damage repair via downregulation of RAD51 has already been reported(*33*). In sensitive cells, a strong lipotoxic phenotype upon combined treatment with palmitate and SSI-4 induced DNA damage as measured by phosphorylated histone H2A.X (Fig.7A). This prompted us to assess the therapeutic potential of SSI-4 combination with the DNA-damaging chemotherapeutic doxorubicin. Indeed, SSI-4 increased doxorubicin induced DNA damage (Fig.7B) with similar effects on lipid peroxidation (Suppl.Fig.8A). Moreover SCD depletion resulted in growth disadvantage in the presence of doxorubicin and increased sensitivity to doxorubicin-induced cytotoxicity (Fig.7C, Suppl.Fig.8B). The relevance of the SFA/MUFA ratio and induction of lipotoxicity in modulating AML cells sensitivity to anthracycline is further supported by the observation that inhibition of palmitate production using a FASN inhibitor or AMPK activation reduces sensitivity to doxorubicin alone or in combination with SSI-4 (Suppl.Fig.8C).

We detected synergism between SSI-4 and doxorubicin in MV-4-11 cells at the majority of dose combinations, while in *iMLL-AF9* cells, which displayed greater sensitivity to doxorubicin, synergy was evident when doxorubicin was applied in lower concentrations (Fig.7D). Moreover, analysis of the BeatAML dataset showed that higher *SCD* expression correlates with reduced sensitivity *in vitro* to cytarabine, an antimetabolite known to cause DNA damage and used in combination with anthracyclines to treat AML patients (Suppl.Fig.8D). Finally, to validate these findings *in vivo* we used two aggressive models of fully established AML that are more representative of the scenario routinely encountered in clinic, a cell line derived xenograft (CDX) and a transplant of leukemic *iMLL-AF9* cells. The latter reached an average leukemic blasts infiltration in PB of 20% before treatment (Suppl.Fig.8E), a common criterion for AML diagnosis. Similarly to what we observed in the PDX, response to SSI-4 therapy as a single agent was varied with significant survival improvement in the CDX model but not in the murine *iMLL-AF9* model. However in both models and consistent with the *in vitro* findings, combining SSI-4 treatment with doxorubicin and cytarabine, a protocol mimicking the standard intensive chemotherapy used in patients, significantly prolonged survival (Fig.7E-F). Together these results demonstrate that SCD inhibition augments efficacy of standard AML chemotherapy likely by enhancing their ability to induce DNA damage.

## DISCUSSION

In this work we sought to understand AML metabolic reliance on FAS to uncover novel therapeutic vulnerabilities. Consistent with observations in several solid cancers(*41–43*), high SCD expression is an adverse prognostic marker in AML. The prognostic role of SCD is likely due to its association with sensitivity to chemotherapy given that biosynthesis of unsaturated fatty acids is enriched in relapsed and chemo-refractory patients. The latter observation emphasizes the potential of SCD inhibition as a treatment strategy in AML particularly when combined with standard chemotherapy as supported by our data. The greatest challenge in targeting SCD has been the lack of available clinical grade inhibitors(*15*). SSI-4 is under clinical development for hepatocellular carcinoma(*5*), and in our study demonstrated potent anti-leukemic effects *in vitro* on a subset of AML samples while showing no general or hematopoietic toxicities.

Although specific mutations can dictate selective metabolic vulnerabilities(*44*), metabolic dependencies can be present across multiple genetic backgrounds as a phenotype bottleneck i.e. a state essential for continued tumorigenesis. This can be advantageous as metabolic vulnerabilities can be exploited in a larger proportion of patients in a mutation-agnostic manner but also highlights the challenge to identify determinants of sensitivity. This is particularly crucial for metabolic inhibitors as they often target pathways central to the function of normal cells/tissues. The favorable toxicity profile of SSI-4 is therefore even more translationally relevant.

Leukemic cells clearly dichotomized in sensitive and resistant to SCD inhibition, a pattern also observed in glioblastoma and melanoma(*6, 45, 46*). AML sensitivity to SSI-4 was not related to mutational background, instead, sensitive cells displayed both greater *de novo* MUFA production and higher MUFA levels. Sensitive cells dependency on FA desaturation caused a greater SFA/MUFA imbalance upon SCD inhibition resulting in lipotoxicity. SCD appears to be the regulatory nexus of *de novo* FAS in AML cells, because oleate decreases FAS both in resistant and sensitive cells, rescuing SSI-4-mediated toxicity both by replenishing the MUFA pool, but also preventing SFA production. Similar effects were observed in pancreatic duct adenocarcinoma (PDAC) cells where exposure to oleate in delipidated medium also decreased FA production, irrespective of SCD inhibition, and inhibition of SFA production rescued toxicity of SCD inhibition, consistent with our model(*47*). It is therefore clear that to drive cytotoxicity via SCD inhibition a significant imbalance between SFA/MUFA needs to be generated. This could be also achieved by modulating the diet, either through a palmitate rich or a caloric restricted diet, which creates a dependency on FAS and reduces SCD levels(*47*) and will be the focus of future work. Conversely, therapeutic interventions which inhibit FAS might reduce the efficacy of SCD inhibition and should be avoided in this setting.

While we did not detect a transcriptional signature of sensitivity, we noted that sensitive primary AML samples displayed increased phosphorylation of IRS2, a direct downstream target of insulin and growth factor receptors(*48*). Insulin is a known regulator of SCD expression(*49*), and sensitivity to SCD inhibition in glioblastoma has been linked to increased ERK phosphorylation(*6*), also a downstream target of RTK signaling(*50*), while FLT3 inhibition in AML decreases MUFA levels through SCD downregulation(*18*). Interestingly, all the sensitive cell lines in our study harbored mutations associated with increased RTK signaling (*BCR-ABL*, *FLT3*) and resistant cell lines presented with *NRAS* mutation, that is known to decrease sensitivity to SCD in solid cancer models(*51*). However, these observations could not be replicated in our primary samples cohort, again highlighting the complexity of interactions between mutational profiles and signaling mechanisms in AML(*52*). Although our data indicate that sensitivity to SCD inhibition might correlate with levels of RTK signaling, further work on larger patient cohorts is required to confirm this as a predictive biomarker of response.

The exact mechanism through which lipotoxicity induces cell death remains ill-defined. A previous report ascribed palmitate-induced toxicity to induction of ER stress(*34*). Conversely, reduction of MUFAs is a known inducer of ferroptotic cell death(*53*). In response to SCD inhibition we observed pleiotropic effects causing both increased ER stress with activation of transcription factor DDIT3/CHOP and apoptotic machinery and elevated lipid peroxidation which is a hallmark of ferroptosis(*32*). However, inhibition of each of these pathways alone could achieve only a partial rescue of SSI-4-mediated cell death, indicating their functional redundancy. Indeed it has already been shown in glioma cells that SCD inhibition results in distinct downstream effects(*28*) and both apoptotic and ferroptotic cell death pathways are triggered in response to SCD inhibition in ovarian cancer(*36*). These conclusions are further supported by the observation that oleate supplementation, which can fully rescue viability of SSI-4 treated cells, acts in parallel on FAS, lipid peroxidation, ER stress and apoptosis, in accordance with its already known ability to rescue both apoptotic and ferroptotic cell death in response to SFA accumulation(*54*).

SSI-4 toxic effects were less pronounced when primary AML cells were grown in co-culture with stroma and were only partially maintained *in vivo*. As expected, and comparable to what is described in glioblastoma(*6*), SSI-4 as a single agent did not consistently prolong survival in AML mouse models. Still, increased lipid peroxidation as a marker of SCD inhibition was maintained even when the decrease in leukemic burden was not prominent, indicating that SSI-4 treated leukemic cells are primed for a second cytotoxic hit. In our models lipotoxicity was linked with increased DNA damage, consistent with what was seen in glioblastoma(*6*), and oxidative stress-inducing strategies have been shown to enhance the response to chemotherapy in AML(*55*). Indeed, we observed synergy between SCD inhibition and doxorubicin *in vitro* and combination of SSI-4 with conventional AML chemotherapy in two aggressive AML models *in vivo* significantly prolonged survival. These findings highlight that genuinely lethal metabolic bottlenecks are often generated by the action of already approved therapeutic interventions and metabolic vulnerabilities can be fully exploited via synergistic combination therapies(*18, 56, 57*). In conclusion our findings support the efforts of devising new treatment approaches in AML focusing on the metabolic axis of MUFA synthesis. Going forward, as will be the case for most metabolic inhibitors, further research on the identification of predictive biomarkers of response and novel combination approaches, with either other approved therapies or dietary interventions, are essential for fully realizing the potential of targeting this axis in AML.

## MATERIALS AND METHODS

### STUDY DESIGN

This study aimed to reveal whether SCD represents a promising therapeutic target in acute myeloid leukemia and determine toxicity and efficacy of clinical grade inhibitor SSI-4 in AML setting. Therefore we used the full spectrum of AML models: patient datasets, cell lines, a cohort of 47 primary AML samples from two independent institutions, murine AML models, cell line- and patient-derived xenograft models. Initial epidemiological studies that attempted to evaluate SCD association with AML prognosis and response to therapy were performed on datasets comprising more than 1000 patients. Cell line studies on a panel of 8 genetically heterogeneous AML cell lines were designed to determine SSI-4 efficiency in inducing anti-leukemic effects, specificity to SCD inhibition, cellular metabolic response and modes of action. A wide cohort of patient samples was screened for translational potential of effects observed on cell lines and identification of potential biomarkers of response using genomic, transcriptomic, proteomic, phosphoproteomic and metabolomic approaches. Mouse models were used to establish general and hematopoietic toxicity of the compound *in vivo* as well as its therapeutic efficacy against AML either alone or in combination with conventional chemotherapy. Animal sample size was determined based on our previous experience and pilot studies. Number of biological replicates is indicated in figure legends and for experimental work involving cell lines, a minimum of three independent experiments were performed. Detailed methods are included in Supplementary methods and all reagents used are listed in Supplementary table 2.

#### Cell culture: cell lines and primary human AML patient derived samples

K-562 (ATCC, CCL-243), MOLM-13, MV-4-11, THP-1, HL-60, Kasumi-1, OCI-AML3, TF-1 (Sanger Institute), 293T-Pheonix cells (kind gift of B. Huntly, University of Cambridge) and MS-5 (DSMZ, ACC 441) cells were cultured following ATCC and DSMZ recommendations. Cell lines used were STR typed and regularly checked for Mycoplasma contamination.

Frozen AML samples from Barts Cancer Institute (n=36) and University Medical Center Groningen (n=11) were retrieved from respective institute’s biobank thawed and plated in co-culture with MS-5 stromal cells. After treatment with SSI-4, viability was determined using anti-Annexin-V antibody in combination with propidium iodide or DAPI stain.

All human samples were obtained and studied after informed consent and protocol approval by both institutions Ethical Committees and BCI Tissue Biobank’s scientific sub-committee in accordance with the Declaration of Helsinki.

#### In vivo experiments

All experiments on animals were performed under UK Home Office authorization. The mice strains used in the study were NBSGW(*58*) and Vav-iCre (*59*) and were purchased from Jackson Laboratory. *iMLL-AF9* mice were described previously(*60*) and were a kind gift of Jürg Schwaller.

Animals were treated orally with 10 or 30 mg/kg SSI-4 in 10% Captisol solution or vehicle control. For experiments involving conventional chemotherapy protocol, it was delivered in a 5 day protocol in which on days 1, 3 and 5 animals intravenously received 1.0 mg/kg doxorubicin and 50 mg/kg cytarabine in the same syringe and on days 2 and 4 animals intravenously received 50 mg/kg cytarabine.

All recipients were culled upon reaching either treatment endpoint or humane endpoint, as noted in survival curves, and their PB, spleen and BM were examined. Complete blood counts and bone marrow cellularity counts were performed using Celltac α Automated Hematology Analyzer (Nihon Kohden).

#### Flow cytometry

Briefly, Cell viability was determined using anti-Annexin V antibody with propidium iodide, 7-AAD or Zombie NIR™ stains. Cell cycle progression was determined using PI solution. Lipid uptake measurement was performed using C1-Bodipy C12 500/510, while lipid peroxidation was measured with Bodipy 581/591 and determined as the ratio of MFI in the green and red channel. Immunophenotypic analyses were performed using antibody cocktails as described in Figure legends and Supplementary methods.

Flow cytometry analyses were performed using LSRFortessa and FACSymphony A3 (BD) instruments and all data analysis was performed using FlowJo 10.0 software.

#### RNA sequencing and analysis

RNA Sequencing and bioinformatics analysis was provided by Novogene UK Company Limited (Cambridge, UK).

#### Glucose labelling

Cells were grown for 24h in RPMI medium with no glucose, supplemented with 10% FBS, 50 IU/ml penicillin and 50 μg/ml streptomycin and 2 g/L U-¹³C_16_-Glucose and treated with SSI-4 (1 µM) with or without the addition of BSA-conjugated sodium oleate (100 µM) or sodium palmitate (100 µM). In analysis, fatty acids containing isotope ¹³C peaks m+0 and m+1 were marked as unlabeled and the ones containing m+2 and higher as labelled.

#### Metabolomics experiments

For lipidomics analysis, lipid species were extracted using monophasic isopropanol extraction and analyzed using liquid chromatography-mass spectrometry following the methodology described before(*61*). Lipid annotation was performed with LipiDex software using the parameters used in the original publication(*62*) and additional analysis of the lipidomics dataset was performed with the LipidSuite webtool (https://suite.lipidr.org).

For fatty acid profiling, apolar metabolites were isolated from cells using chloroform:methanol extraction and fatty acids partitioned from polar metabolites by resuspension of dried extracts in chloroform:methanol:water. Data acquisition was performed using gas chromatography – mass spectrometry. Data was acquired using MassHunter software (version B.07.02.1938). Data analysis was performed using MANIC software, an in house-developed adaptation of the GAVIN package(*63*). Fatty acids were identified and quantified by comparison to authentic standards and ^13^C_1_-lauric acid as an internal standard.

### STATISTICS AND REPRODUCIBILITY

All data was analyzed and visualized in Prism 9.0 (GraphPad) and all data are shown as mean ± standard error of the mean, unless otherwise stated. The cohorts were dichotomized into groups with high and low gene expression after calculating the optimal cut point value using the receiver operating characteristic (ROC) curve for censored overall survival data. Overall survival was plotted using Kaplan–Meier plots, using Cox proportional hazard regression to compare the differences between the curves, providing the Hazard ratio (HR) and the 95% confidence interval (CI). According to data availability, we adjusted prognosis prediction for confounders as follows: age (as continuous variable), sex (male vs. female), white blood cell counts (WBC, as continuous), and European LeukemiaNet categorization (ELN2010 or ELN2017). The difference between multiple experimental groups was analyzed by two-way ANOVA (Kruskal–Wallis test, post hoc Dunn analysis) or two-tailed paired or unpaired Student’s t test. IC50 and cell viability curves were determined using non-linear regression analyses. Correlation was calculated using Pearson or Spearman correlation coefficients. P-values are indicated in Figure legends.

## List of Supplementary Materials

Materials and Methods

Fig. S1 to S8

Tables S1 to S2 for multiple supplementary tables

Data files S1 to S3 (Excel files)

## Acknowledgments

We would like to thank Dora Visnjic for helpful advice and discussion and Han Tun and Justyna Gleba for critically reading the manuscript. We wish to thank the Barts Cancer Institute tissue bank for sample collection and processing.

## Funding

Cancer Research UK Advanced Clinician Scientist fellowship C57799/A27964 (PG) The Lady Tata Memorial Trust International Award for Research in Leukaemia (VD) Cancer Research UK Core Award C16420/A18066 (BCI Flow cytometry facility)

## Author contributions

Conceptualization: PG, VD.

Methodology: VD, HL, RB, CP, MA, GRB, AvK, BP, JES, JT, AF, JAC, BP (2), KK,

PG. Validation: LO.

Formal analysis: ARM, PCI, FBC, DPM, GRB, EB.

Investigation: VD, HL, RB, CP, SJ, SA, JD, LW, JC, AM, KW, MA, GRB, LO.

Resources: CP, JES, JS, JT, JJS, LH, JAC, MY, PC, JF, KRP, KK, PG.

Writing – original draft: VD, PG.

Writing – review and editing: all authors.

Visualization: VD, PG.

Supervision: KRP, KK, PG.

Project administration: PG.

Funding acquisition: AvK, PC, JF, KK, PG.

## Competing interests

LH is CEO and co-founder of Modulation Therapeutics, the company holding intellectual property to SSI-4. J.A.C. holds a patent regarding use of the SCD inhibitor SSI-4.

## Data and materials availability

All regents and materials in this study are listed in Supplementary table 1. SSI-4 can be obtained through material transfer agreement (MTA) with Modulation therapeutics Inc. All cell lines generated in this study can be obtained upon request.

The RNA-sequencing data generated in this study have been deposited in the ArrayExpress database and are available at E-MTAB-13174. Data from free fatty acid profiling and lipidomics analyses are available in Supplemental data 1, 2 and 3.

DNA sequencing, RNA sequencing, proteome and phosphoproteome data on BCI primary AML samples were derived from previously published studies from Barts Cancer Institute(*52, 64*).

Publicly available clinical and transcriptomic data of five adult AML cohorts whose patients were treated with intensive chemotherapy were used to investigate the prognostic role of SCD expression: AML TCGA (data obtained from https://www.cbioportal.org/), GSE6891, GSE425 (Bullinger), GSE10358, GSE14468. Normalized gene expression data were retrieved from the Gene Expression Omnibus (GEO) database (www.ncbi.nlm.nih.gov/geo/). SCD signature was generated by dichotomizing patients in TCGA dataset in high and low expressing samples based on median expression. Dataset GSE97631 was used to determine SCD signature expression in MRD stage.

Data from manuscript 10.1056/NEJMoa1808777 (NEJM1808777),(*65*) and GSE66525 were used to perform comparative RNASeq analyses on paired diagnosis-relapse samples in human cohorts and murine models.

SCD signature was generated by dichotomizing patients in TCGA dataset in high and low expressing samples based on median expression. Dataset GSE97631 was used to determine SCD signature expression in MRD stage.

BeatAML dataset (data obtained http://www.vizome.org/), was used to determine association of SCD expression with sensitivity to cytarabine *ex vivo*.

Data from manuscript 10.1016/j.cell.2019.12.023 were used for proteomics comparison of sensitive and resistant cell lines. Normalization method used is described in the manuscript(*25*). All code and data analyses are available upon request.

## Notes

### Summary of Updates

Minor textual updates for clarity, Figure 6 revised, ORCID links added

