## Supplemental methods, figures and tables for "SCD inhibition preferentially eradicates AML displaying high de novo fatty acid desaturation and synergizes with chemotherapy"

*Cell culture*

K-562 (ATCC, CCL-243), MOLM-13, MV-4-11, THP-1, HL-60, Kasumi-1, OCI-AML3, TF-1 (Sanger Institute), 293T-Pheonix cells (kind gift of B. Huntly, University of Cambridge) and MS-5 (DSMZ, ACC 441) cells were cultured following ATCC and DSMZ recommendations. Cell lines used were STR typed and regularly checked for Mycoplasma contamination.

For the purpose of exogenous addition of fatty acids, sodium oleate (Sigma-Aldrich) was dissolved in sterile water and conjugated with fatty acids-free bovine serum albumin (Sigma-Aldrich) in ratio 1:2 at 37 °C for 20 minutes. Sodium palmitate (Sigma-Aldrich) was dissolved in 100% ethanol at 75 °C for 10 minutes and conjugated with fatty acids-free bovine serum albumin in ratio 1:3 at 37 °C for 20 minutes. Dissolved sodium oleate was stored at -20°C, and dissolved sodium palmitate was prepared fresh.

*Primary human AML patient derived samples*

Frozen AML samples (n=36) were retrieved from Barts Cancer Institute Biobank. Upon thawing, T cells were depleted using EasySep™ Human TCR Alpha/Beta Depletion Kit (Stem Cell Technologies). Enriched samples were plated in concentration 0.4 – 1.0 X 106/mL in Myelocult H5100 medium (Stem Cell Technologies) supplemented with 20 ng/mL IL-3, G-CSF and TPO (Biolegend) either in liquid culture or in co-culture with irradiated MS-5 cells and treated with SSI-4 (1 µM) or vehicle control with or without addition of palmitate (100) or vehicle control for 7 days. After 3 days of treatment, half of medium was exchanged with fresh medium containing the corresponding agent. After 7 days, viability was assessed using Annexin V FITC/PI stain and cell cycle was measured using PI solution. Samples were deemed sensitive to treatment if a decrease in viability greater than 5% was detected.

Samples from University Medical Center Groningen (UMCG) (n=11) were thawed and plated in co-culture with MS-5 stromal cells in Gardner’s medium with the addition of G-CSF, IL3, romiplostim, SR1 and UM171. After 2 days recovery, SSI-4 was added in two concentrations (1 and 10 µM). After 1 and 4 days of treatment, cell viability was assessed using MACSquant flow cytometer (Miltenyi Biotec), viable cells were determined as Annexin-V^-^/DAPI^-^, and area under curve (AUC) for drug sensitivity was calculated.

*In vivo experiments*

All experiments on animals were performed under UK Home Office authorisation. The mice strains used in the study were C57BL/6, NOD.Cg-*Kit^W-41J^ Tyr* ^+^ *Prkdc^scid^ Il2rg^tm1Wjl^*/ThomJ (NBSGW) and Vav-iCre and all were purchased from Jackson Laboratory. *iMLL-AF9* mice were a kind gift of Jürg Schwaller. All transgenic and knockout mice were CD45.2^+^. Congenic recipient mice were CD45.1^+^/CD45.2^+^. Mice used for support BM cells during transplantation experiments were CD45.1^+^.

For SSI-4 toxicity experiment, 13- to 15-week-old mixed gender C57BL/6 mice were treated with 10 or 30 mg/kg SSI-4 in 10% Captisol solution or vehicle control orally. All recipients were culled upon reaching treatment endpoint and their PB, spleen and BM were examined for SSI-4 effects on normal haematopoiesis. Complete blood counts and bone marrow cellularity counts were performed using Celltac α Automated Hematology Analyzer (Nihon Kohden).

For syngeneic leukaemia model, CD45.1^+^/CD45.2^+^ recipient mice were lethally irradiated using a split dose of 8 Gy (two doses of 4 Gy administered at least 4 hours apart) at an average rate of 1.086 Gy/min using a RADSOURCE X-ray irradiator. 2,000 *iMLL-AF9* LSK cells were transplanted into lethally irradiated CD45.1^+^/CD45.2^+^ recipient mice together with 200,000 unfractionated support CD45.1^+^ wild-type BM cells. Engraftment and leukemic burden in PB was assessed three weeks after transplantation and animals were treated orally with 10 mg/kg SSI-4 in 10% Captisol solution or vehicle control.

*iMLL-AF9* mice were treated with doxycycline to initiate leukemic transformation. LSK cells from these mice were grown in a CFC assay in MethoCult M3434 supplemented with 250 ng/mL doxycycline for 6 days to establish their colony forming potential. 5,000 transformed leukemic cells were tail vein injected into non-irradiated 8- to 13-week-old mixed gender NBSGW mice. When leukemic burden in PB reached on average 20%, animals were randomized into treatment groups and subjected to combined chemotherapy and SSI-4 treatment. Chemotherapy was delivered in a 5 day protocol in which on days 1, 3 and 5 animals intravenously received 1.0 mg/kg doxorubicin and 50 mg/kg cytarabine in the same syringe and on days 2 and 4 animals intravenously received 50 mg/kg cytarabine. In parallel with chemotherapy, SSI-4 was delivered intraorally in the dose 10 mg/kg.

For xenotransplantation experiments, 100,000 MV-4-11 cells were tail vein injected into non-irradiated 10- to 12-week-old mixed gender NBSGW mice and began drug treatment 14 days after transplantation. Chemotherapy protocol was delivered as previously described and SSI-4 was delivered intraorally in the dose 10 mg/kg.

For patient derived xenografts (PDX), patient samples AML3 and AML5 were T-cell depleted and tail vein injected into non-irradiated 8- to 10-week-old mixed gender NBSGW mice (2 and 4 million cells per mouse, respectively). Engraftment of human hematopoietic cells was confirmed at weeks 8 and 10 by BM sampling and testing for the presence of human CD45^+^ cells. Upon engraftment confirmation, mice were randomized per treatment group and orally treated with 10 mg/kg SSI-4 in 10% Captisol solution or vehicle control.

For doxycycline (Dox) treatment needed for leukemic transformation of *iMLL-AF9* model, mice were provided *ad libitum* access to drinking water containing 2 mg/mL DOX with 30% sucrose. All animals were culled upon reaching either treatment endpoint or their humane endpoint as recorded in survival curves.

*Leukaemic transformation*

*Meis1/Hoxa9* transformed murine leukemic cells and *iMLL-AF9* murine leukemic cells were grown in IMDM supplemented with 10% FBS and 10 ng/ml SCF, 5 ng/ml IL-3 and 5 ng/ml IL-6 or 20 ng/ml SCF, 10 ng/ml IL-3, 10 ng/ml IL-6 and 250 ng/mL doxycycline respectively.

*Flow cytometry*

Cell viability was determined using FITC-conjugated anti-Annexin V antibody with propidium iodide or Zombie NIR™ (BioLegend) stains or PE-conjugated anti-Annexin V antibody with 7-AAD stain. Double negative cells were deemed viable. Cell cycle progression was determined using PI solution (50 μg/ml PI, 10 mm Tris, pH 8.0, 10 mm NaCl, 10 μg/ml RNase A, 0.1% Igepal) and percentage of cells in each cell cycle phase was calculated using Dean Jett Fox model.

For lipid peroxidation measurement cells were incubated with 4 µM Bodipy 581/591 for 30 min at 37 °C with gentle shaking, protected from light. Cells were washed twice with PBS and analysed by flow cytometry. Lipid peroxidation was determined as the ratio of MFI in the green channel (530) vs MFI in the red channel (610) per manufacturer’s instructions.

For lipid uptake measurement cells were incubated with 1 µM C1-Bodipy C12 500/510 at 37 °C and 5% CO_2_ for 24 h. Cells were washed twice with PBS and analysed by flow cytometry.

BM cells were isolated by crushing tibias and femurs using a pestle and mortar. Splenic cells were prepared by mashing the tissue and passing through a 70 µm strainer. Erythrocytes in PB were lysed using ammonium chloride solution. Single cell suspensions from BM, spleen or PB were incubated with Fc block and then stained with antibodies. For HSC and progenitor cell analyses, unfractionated BM cells were stained with lineage markers containing biotin-conjugated anti-CD4, anti-CD5, anti-CD8a, anti-CD11b, anti-B220, anti-Gr-1 and anti-Ter119 antibodies together with BV711-conjugated anti-c-Kit, APC-Cy7-conjugated anti-Sca-1, PE-conjugated anti-CD48 and PE-Cy7-conjugated anti-CD150 antibodies. Biotin-conjugated antibodies were then stained with PB-conjugated streptavidin. For analyses of differentiated cells, PB was stained with PerCP-conjugated anti-B220 and APC-Cy7-conjugated anti-CD19 antibodies for B cells; APC-conjugated anti-CD11b and PE-Cy7-conjugated anti-Gr-1 for myeloid cells; PE-conjugated anti-CD4 and anti-CD8 antibodies for T cells.

To distinguish CD45.2^+^-donor derived cells in PB or BM of transplanted mice, BV711-conjugated anti-CD45.1 and FITC-conjugated anti-CD45.2 antibodies were used, and to distinguish human versus mouse CD45^+^ positive cells PB-conjugated anti-human CD45 and APC-conjugated anti-mouse CD45 antibodies were used. TO-PRO-3 was used for dead cell exclusion. Human myeloid cells in xenograft models were distinguished using PE-conjugated anti-human CD33 and B-cell exclusion was done using BV711-conjugated anti-human CD19.

Flow cytometry analyses were performed using LSRFortessa and FACSymphony A3 (BD) instruments and all data analysis was performed using FlowJo 10.0 software.

*Lentiviral transduction*

293T-Pheonix cells were transfected with pMD2.G, psPAX2 and construct plasmid using TransIT (Mirus Bio) for transient transfection. After overnight incubation at 37 °C with 5% CO2, media containing virus was harvested and filtered through a 0.45 µm filter to remove any cells. For transduction of cell lines, 1.8 million cells were spinoculated with virus containing medium and 4 µg/ml polybrene (Santa Cruz Biotechnology, TR1003) for 90 min at 900 × g. After 24 h, free viral particles removed by washing three times with sterile PBS and cells were subjected to 2 µg/mL puromycin selection for 14 days.

*Generation of CRISPR knockout clones*

The knock-out of genes was accomplished using the CRISPR/Cas9 system. For this MV411 and THP-1 cell lines were transduced with Cas9 lentivirus (Addgene, lentiCas9-Blast), generating Cas9 expressing cell lines. Functional gRNA sequences (Supplemental table 1) were obtained from genome-wide gRNA library and ligated in the backbone, conjugating SCD gRNA with BFP (Addgene, pKLV2-U6gRNA5(BbsI)-PGKpuro2ABFP-W). Virus for Cas9, SCD gRNA and NT gRNA were produced and transduced as described above. Obtaining single cell colonies was performed using methylcellulose media (H4531; STEMCELL Technologies, Cambridge, UK).

*Western blotting*

Cells were lysed using cell lysis buffer (Cell Signaling Technology) supplemented with protease and phosphatase inhibitors (Sigma-Aldrich, Millipore). DNA was sheared by passing the sample through a fine gauge needle 8 times before centrifugation for 10 min at 14,000 x g. The protein extracts were subjected to SDS–PAGE (Invitrogen) and then transferred onto PVDF membranes (Sigma-Aldrich) using wet transfer. Membranes were blocked with 5% milk powder in 1x PBS-T for 30 mins then incubated overnight at 4 °C with primary antibodies. After 2h incubation at RT with appropriate horseradish peroxidase-coupled secondary antibody, bands were detected with enhanced chemiluminescence substrates (BioRad, ThermoFisher) and visualized using an Amersham Imager (GE Healthcare) or a ChemiDoc Imaging system (BioRad). For analysis of phosphorylated vs total proteins equal aliquots of protein solution were loaded to parallel gels and processed simultaneously. Band intensity quantification (densitometry) was performed using Image J software and normalized to ß-actin. Antibodies used are enclosed in Supplemental table 1.

*Real Time quantitative PCR*

The RNA was isolated using Direct-zol RNA Miniprep kit (Zymo Research) following the manufacturer’s protocol and reverse transcribed using High-Capacity cDNA Reverse Transcription Kit (Applied Biosystems). For real-time quantitative PCR (qPCR), 5 ng of cDNA, 5 µL of PowerUp SYBR Green MasterMix (Applied Biosystems) and 2 pmol of primers (Suppl. table 2) were used per well of 384-well plate. Reactions were performed in triplicate using C1000 Thermocycler 384well (BioRad). Gene expression was quantified using comparative ΔΔ-Ct method and *ACTB* was used as the housekeeping gene. Data is expressed as log2 fold change in comparison to control sample and represents results of three independent experiments measured in duplicate.

*RNA sequencing and analysis*

RNA Sequencing was provided by Novogene UK Company Limited (Cambridge, UK). The RNA was isolated using Direct-zol RNA Miniprep kit (Zymo Research) following the manufacturer’s protocol. Messenger RNA was purified from total RNA using poly-T oligo-attached magnetic beads (poly-A enrichment). Generated library was sequenced on an Illumina platform and paired-end reads were generated.

Reads were mapped to a reference genome using Hisat2 and Featurecounts was used to count the read numbers mapped of each gene. Differential expression analysis of two conditions was performed using the edgeR R package. Corrected p-value of 0.005 and |log2(Fold Change)| of 1 were set as the threshold for significantly differential expression. Enrichment analysis based on KEGG pathways (<http://www.genome.jp/kegg/>) was performed using clusterProfiler R package or Enrichr webtool (<https://maayanlab.cloud/Enrichr/>). Gene set enrichment analyses (GSEA) and single sample GSEA (ssGSEA) were performed using the local version of the GSEA analysis tool (http://www.broadinstitute.org/gsea/index.jsp). For GSEA analysis normalized read counts for all conditions were used and genes were ranked using the signal-to-noise metric and FDR and NES were calculated using 1000 gene-set permutation.

*Glucose labelling*

Cells were grown for 24h in RPMI medium with no glucose, supplemented with 10% FBS, 50 IU/ml penicillin and 50 μg/ml streptomycin and 2 g/L U-¹³C_16_-Glucose (CK Isotopes) and treated with SSI-4 (1 µM) with or without the addition of BSA-conjugated sodium oleate (100 µM) or sodium palmitate (100 µM). At the end of the treatment, cells were counted in triplicates for normalization, washed in cold PBS and apolar fraction of cell pellets was isolated using methanol:chloroform extraction. In analysis, fatty acids containing isotope ¹³C peaks m+0 and m+1 were marked as unlabelled and the ones containing m+2 and higher as labelled.

*Metabolomics experiments*

For lipidomics analysis, lipid species were extracted from cell pellets using monophasic isopropanol extraction and analysed using liquid chromatography and a Q Exactive™ Hybrid Quadrupole-Orbitrap™ Mass Spectrometer (ThermoFisher). Peak detection, alignment and deconvolution was performed with Compound Discoverer software (ThermoFisher) and lipid annotation was performed with LipiDex software. Additional analysis of the lipidomics dataset was performed with the LipidSuite webtool (https://suite.lipidr.org).

For fatty acid profiling, apolar metabolites were isolated from cells using chloroform:methanol extraction (2:1, v/v, both HPLC grade, Fisher) and fatty acids partitioned from polar metabolites by resuspension of dried extracts in chloroform:methanol:water (1:3:3, v/v, HPLC grade, Fisher). Data acquisition was performed using an Agilent 7890B-7000C GC-triple-quadrupole MS in EI mode after derivatization of twice methanol-washed dried lower (apolar) phase by addition of 25 μL chloroform/methanol (2:1, v/v) and 5 μL tetramethylammonium hydroxide (TMAH, RT, no incubation) for fatty acids. GC-MS parameters were as follows: carrier gas, helium; flow rate, 0.9 mL/min; column, DB-5MS (Agilent); inlet, 250°C; temperature gradient, 70°C (1 min), ramp to 230°C (15°C/min, 2 min hold), ramp to 325°C (25°C/min, 3 min hold). Scan range was m/z 50-565. Data was acquired using MassHunter software (version B.07.02.1938). Data analysis was performed using MANIC software, an in house-developed adaptation of the GAVIN package. Fatty acids were identified and quantified by comparison to authentic standards and ^13^C_1_-lauric acid as an internal standard (Cambridge Isotope Laboratories).

**Supplemental Fig. 1. Increased biosynthesis of unsaturated fatty acids correlates with worse prognosis and relapse in AML.**

(A) Kaplan-Meier curve comparing overall survival in BCI AML patients cohort with adverse prognosis after ELN (n=54) dichotomized using median SCD expression. Log rank (Mantel-Cox) test was used for determining significance. (B) Significantly enriched KEGG signatures in patients with highest SCD expression (n=10) compared with patients with lowest *SCD* expression (n=10) in BCI AML patients cohort with adverse prognosis after ELN (n=54) ranked by combined score from Enrichr enrichment analysis. (C) Gene set enrichment analysis (GSEA) for KEGG pathway Biosynthesis of unsaturated fatty acids in paired relapse-diagnosis samples (GSE66525). (D) Cell cycle analysis of MOLM-13 and MV-4-11 cells after 24h of SSI-4 (1 µM) treatment. (E) Cytogenetic and genetic characteristics of SSI-4 –sensitive and –resistant AML cell lines used in this study.

**Supplemental Fig. 2. Sensitivity to SSI-4 correlates to sensitivity to other means of SCD inhibition.**

(A) MOLM-13, MV-4-11, OCI-AML3, THP-1 and HL-60 cells were treated with A939572 (10 nM, 100 nM, 1 µM) or corresponding vehicle for 72h. Cells with less than 10% decrease in viability were designated to the resistant group. Results are presented as non-linear regression of normalized response. (B) Western blot confirming SCD downregulation in MV-4-11 and THP-1 cells transduced with NT gRNA, SCD gRNA1 and SCD gRNA2. Competition growth assays measuring the ratio of BFP -positive (gRNA expressing cells) to BFP-negative cells (wild type) after 3 and 5 days in MV-4-11 and THP-1 cell lines. (C) Western blot showing the response to 48h of hypoxia (3% O_2_) in MV-4-11 cells with downregulated SCD. In parallel experiments, percentage of viable MV-4-11 cells with downregulated SCD was determined after 72h of hypoxic conditions. (D) Percentage of viable THP-1 cells with downregulated SCD after 72h of hypoxic conditions. (E) K562, MOLM-13, MV-4-11, OCI-AML3, THP-1, HL-60 and Kasumi-1 cells were treated with SSI-4 (1 µM) and lipid uptake was measured using Bodipy 500/510 stain as well as the expression of LDLR and CD36. Red bars present samples where difference between SSI-4 treated and control cells was statistically significant. Viable cells were determined as Annexin-V^-^/PI^-^.* p < 0.05, ** p < 0.01, *** p < 0.001,

**Supplemental Fig. 3. Fatty acids profiles in SSI-4 sensitive and resistant cells in the presence of oleate and palmitate.**

(A) Levels of ^13^C-glucose labeled (m+2 and higher) and unlabeled (m+0, m+1) saturated fatty acids (SFA) palmitate (C16:0) and stearate (C18:0) in MOLM-13, MV-4-11 and OCI-AML3 cells after 24h of SSI-4 (1 µM) treatment with or without the addition of oleate (100 µM) or palmitate (100 µM). (B) Levels of ^13^C-glucose labeled (m+2 and higher) and unlabeled (m+0, m+1) monounsaturated fatty acids (MUFA) palmitoleate (C16:1) and oleate (C18:1) in MOLM-13, MV-4-11 and OCI-AML3 cells after 24h of SSI-4 (1 µM) treatment with or without the addition of oleate (100 µM) or palmitate (100 µM). (C) Ratio of C16 and C18 SFA and MUFA normalized to control in corresponding experimental conditions.

**Supplemental Fig. 4. Sensitivity to SCD inhibition correlates with the activity of fatty acid synthesis pathway.**

(A-B) Western blot analysis of ACC levels and SREBP1 cleavage in response to SSI-4 (1 µM, 24h) upon co-treatment with either oleate (100 µM) or MK-8722 (10 µM). (C) MOLM-13 cells were treated for 72h with SSI-4 (1 µM) with or without addition of ACC inhibitor PF-05221304 (5 µM) or AMPK activator metformin (1 and 5 mM). (D) DepMap dataset was analyzed for correlation (Pearson r) between SCD Crispr scores and expression levels of *ACACA (*coding for ACC*)* and *FASN.* Results on all cancer cell lines in dataset are presented in upper panels and results on AML cell lines in lower panels. Viable cells were determined as Annexin-V^-^/PI^-^.* p < 0.05, ** p < 0.01, *** p < 0.001, **** p < 0.0001.

**Supplemental Fig. 5. SSI-4 induces both lipid peroxidation/ferroptotic cell death and apoptosis.**

(A) RNA sequencing results for MV-4-11 cells treated with SSI-4 (1 µM) for 24h with (right panel) or without the presence of oleate (100 µM) (left panel). (B) Lipid peroxidation measured by Bodipy C11 in MV-4-11 cells with downregulated SCD after 72h in hypoxic conditions (3% O_2_). Representative western blot (n=3) demonstrating SCD levels in normoxic and hypoxic conditions is shown. (C) Lipidomics analysis on MV-4-11 cells treated with SSI-4 (1 µM) for 24h with the presence of oleate (100 µM). Graph represents enrichment analysis per lipid groups of SSI-4 + oleate treated cells in comparison to oleate alone (Q1-Q3 with line at median value) with statistically significant lipid groups marked in red. (D-E) Rescue of lipid peroxidation induction in response to SSI-4 (1 µM, 24h) using oleate (100 µM), as well as lipid peroxidation inhibitors ferrostatin-1 (5 µM) and liproxstatin (2 µM) in MV-4-11 and MOLM-13 cells. Lipid peroxidation was measured was measured using Bodipy C11. (F) MV-4-11 and MOLM-13 cells were treated with SSI-4 (1 µM) with or without addition of liproxstatin (2 µM) for 72h. (G) K562 cells were treated for 72h with SSI-4 (1 µM) with or without addition of ferrostatin-1 (5 µM). Viable cells were determined as Annexin-V^-^/PI^-^. * p < 0.05, ** p < 0.01, *** p < 0.001, **** p < 0.0001.

**Supplemental Fig. 6. SCD inhibition results in broad activation of ER stress response**

(A) MOLM-13 and MV-4-11 cells were treated for 24h with SSI-4 (1 µM) with or without addition of oleate (100 µM) and expression of ER Stress related genes was determined using qPCR. Heatmap represents Log_2_ fold changes of genes measured normalized to ß-actin and control sample (2^-ΔΔCt^). (B) MOLM-13 cells were treated for 72h with SSI-4 (1 µM) with or without addition of PERK inhibitor GSK2656157 (5 µM) IRE-1 inhibitor 4µ8c (10 and 20 µM) and ATF6 inhibitor Ceapin A7 (5 µM) Viable cells were determined as Annexin-V^-^/PI^-^. * p < 0.05, ** p < 0.01, *** p < 0.001, **** p < 0.0001.

**Supplemental Fig. 7. Stromal microenvironment decreases sensitivity to SCD inhibition in primary AML samples characterized by increased signaling downstream of receptor tyrosine kinases.**

(A) Primary AML samples sensitive to SCD inhibition (n=3) were treated with SSI-4 (1 µM) in parallel in liquid culture or co-culture with stroma for 7 days. (B) Primary AML samples were treated with SSI-4 (1 µM) with or without addition of palmitate (1 µM) in liquid culture (n=8) or co-culture with stroma (n=15) for 7 days. (C) A separate AML patients cohort from University of Groningen Medical Center (n=11) was treated with SSI-4 (1 and 10 µM) in co-culture with stroma for 4 days and sensitivity to SSI-4 was expressed as area under curve (AUC). (D) Phosphoproteomic analysis of 5 sensitive and 9 resistant AML patients from BCI Adverse prognosis cohort together with an independent phosphoproteomic analysis (Leukemia 2018) of 3 sensitive and 5 resistant AML patients. Heatmap represents log_2_ fold change of phosphorylated sites on IRS2 in sensitive and resistant samples normalized on target relative intensity. (E) Phosphoproteomic analysis for phosphorylated AKT2 and PLEKHG3 in 5 sensitive and 9 resistant AML patients from BCI Adverse prognosis cohort expressed as log_2_ fold change towards average target relative intensity in analysis. (F) Cell cycle analysis of sensitive (n=7) and resistant (n=8) primary AML samples. (G) Percentage of human CD45^+^ cells in bone marrow of mice in PDX AML models before and after treatment with SSI-4. In the time point before SSI-4 treatment bone marrow was obtained by bone marrow aspiration and based on engraftment levels animals were assigned to comparable control and treatment groups. Viable cells were determined as Annexin-V^-^/PI^-^ and normalized to control. * p < 0.05, ** p < 0.01, *** p < 0.001, **** p < 0.0001.

**Supplemental Fig. 8. Lipotoxicity enhances the anti-leukemic effects of doxorubicin.**

(A) Lipid peroxidation in MV-4-11 cells treated for 72h with SSI-4 (1 µM) with or without addition of doxorubicin (1 µM) was measured using Bodipy C11. (B) Competition growth assay between BFP^+^ and BFP^-^ MV-4-11 cells (MV-4-11 WT *Cas9* expressing) for cells expressing NT gRNA, SCD gRNA 1 and SCD gRNA 2 in the presence of doxorubicin (1 µM). Values are normalized to NT gRNA and expression levels of SCD in all treated groups are shown in representative western blot. (C) MV-4-11 cells were treated for 72h with SSI-4 (1 µM) with or without addition of doxorubicin (1 µM) and FASN inhibitor Fasnall (20 µM) or AMPK activator MK-8722 (10 µM). (D) Correlation of SCD expression and sensitivity to cytarabine in samples from BeatAML 2.0 dataset. (E) Percentage of CD45.2+ leukemia blasts in peripheral blood of NBSGW mice before starting treatment. Viable cells were determined as Annexin-V^-^/PI^-^. * p < 0.05, ** p < 0.01, *** p < 0.001, **** p < 0.0001.

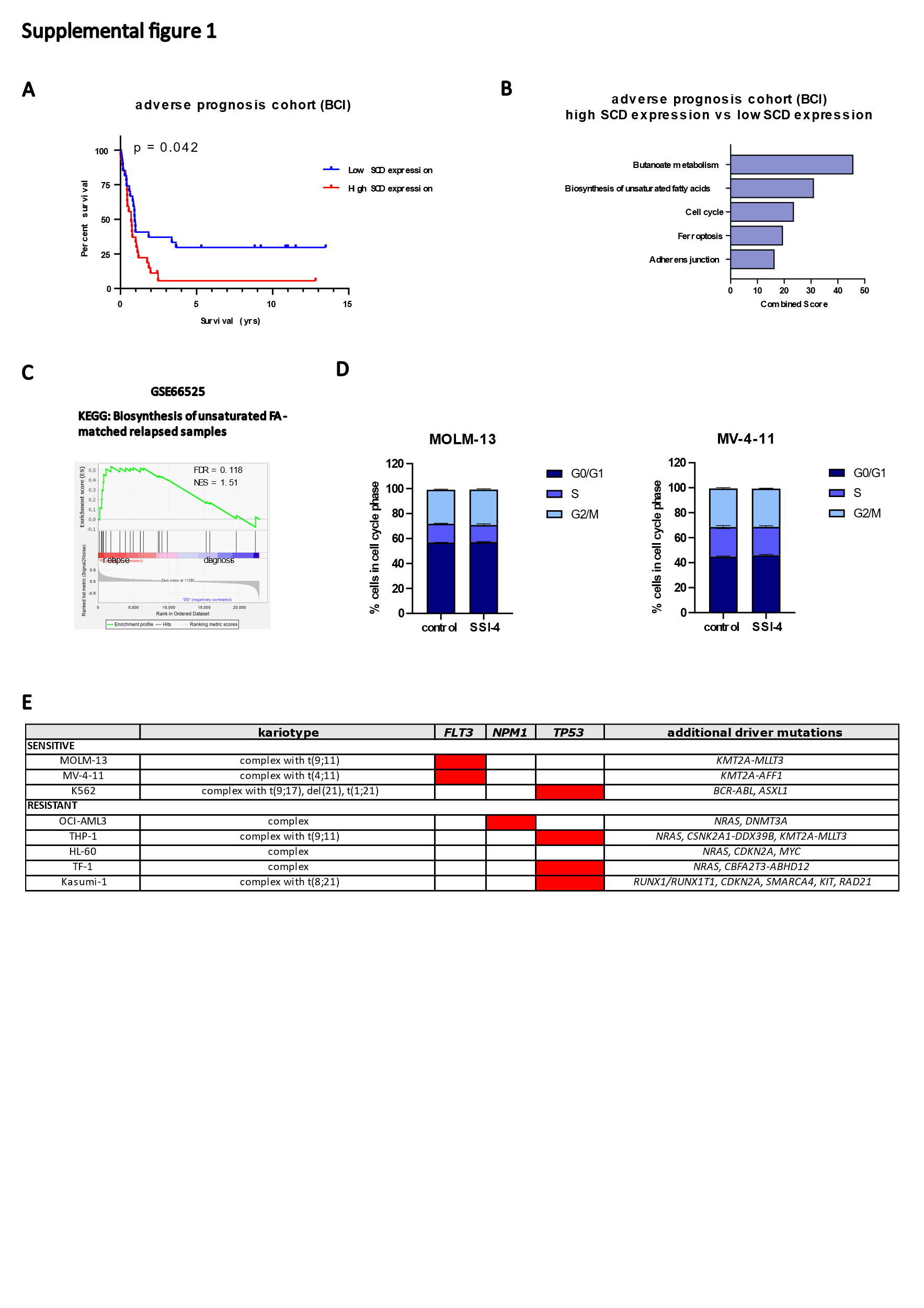

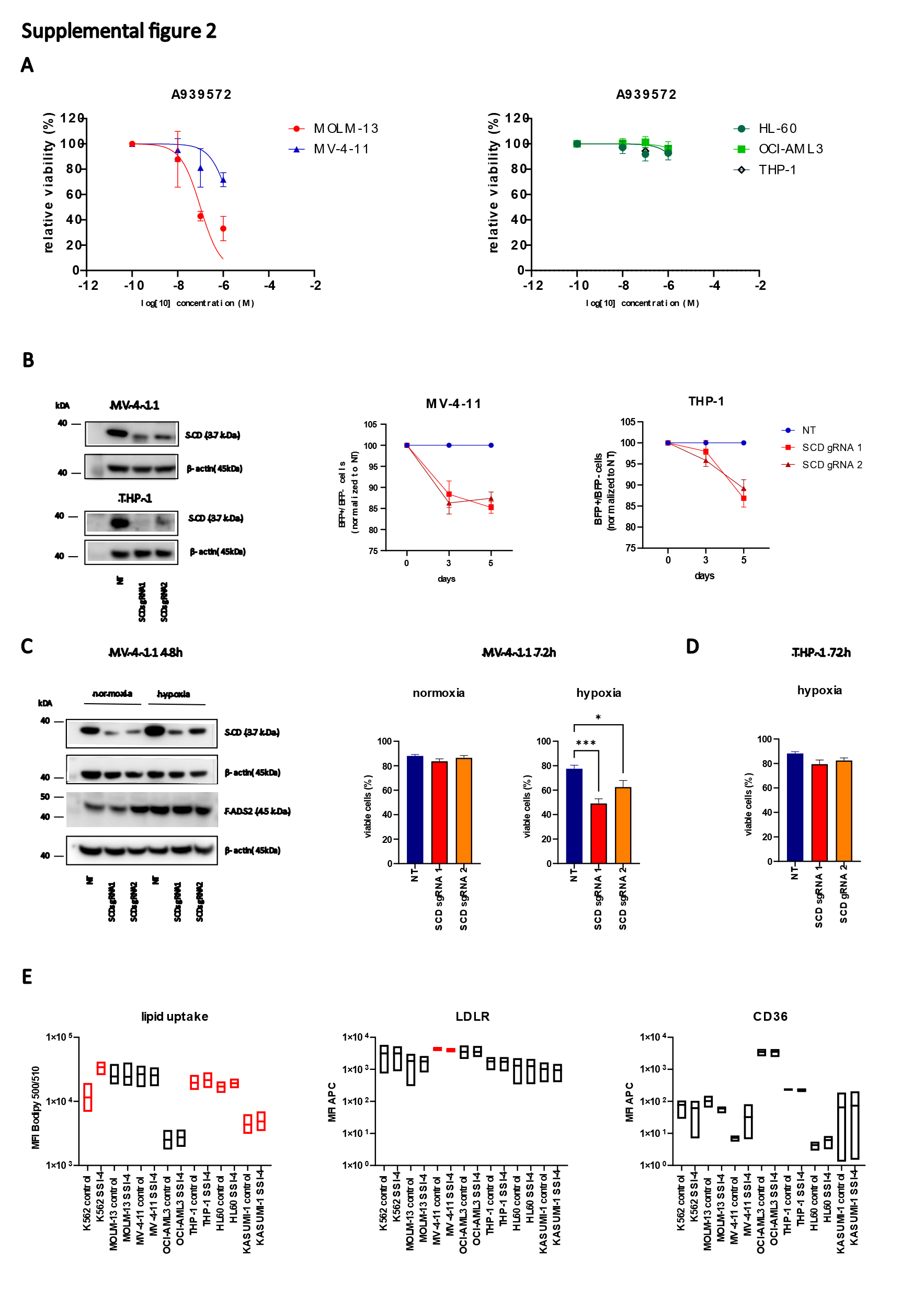

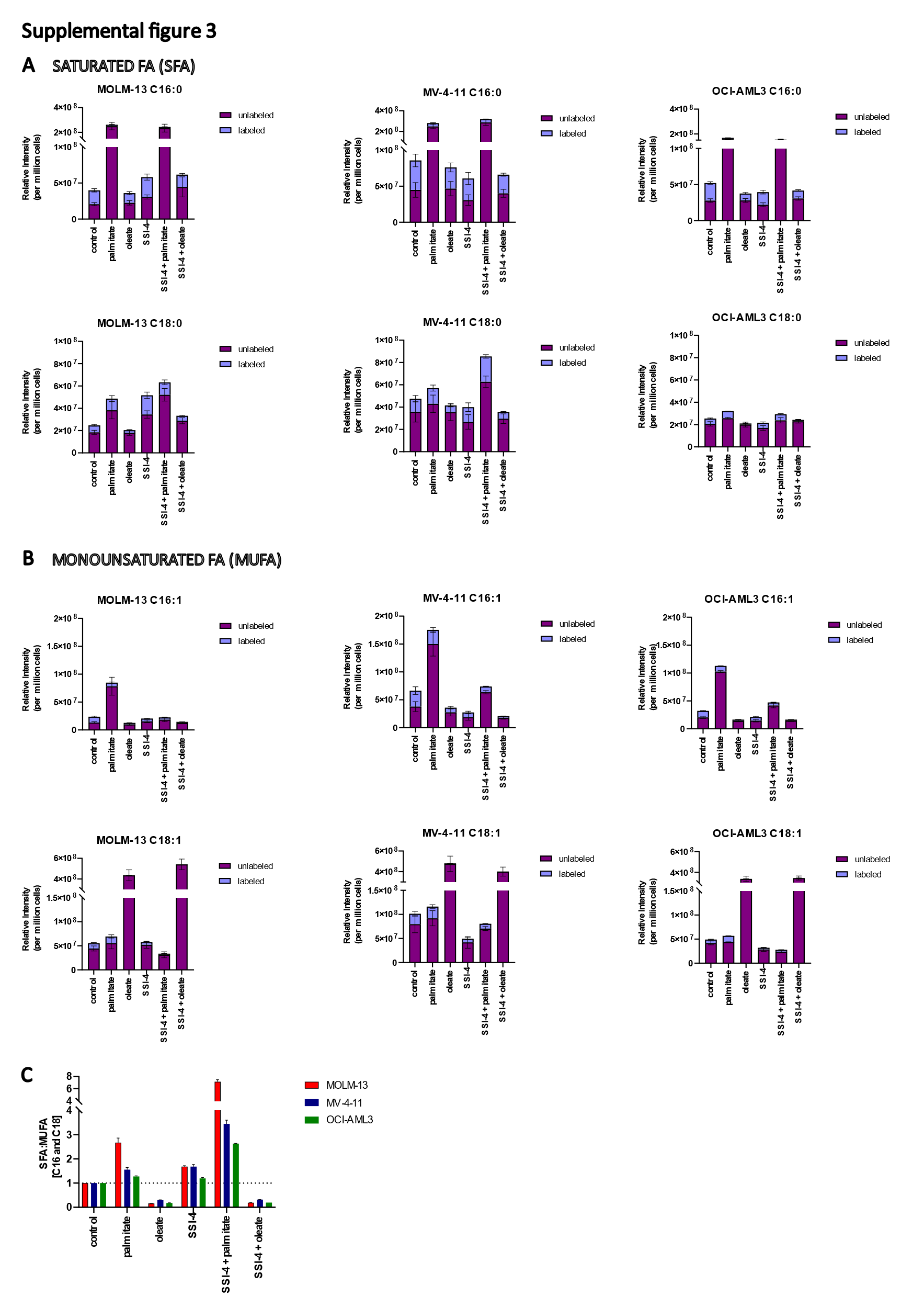

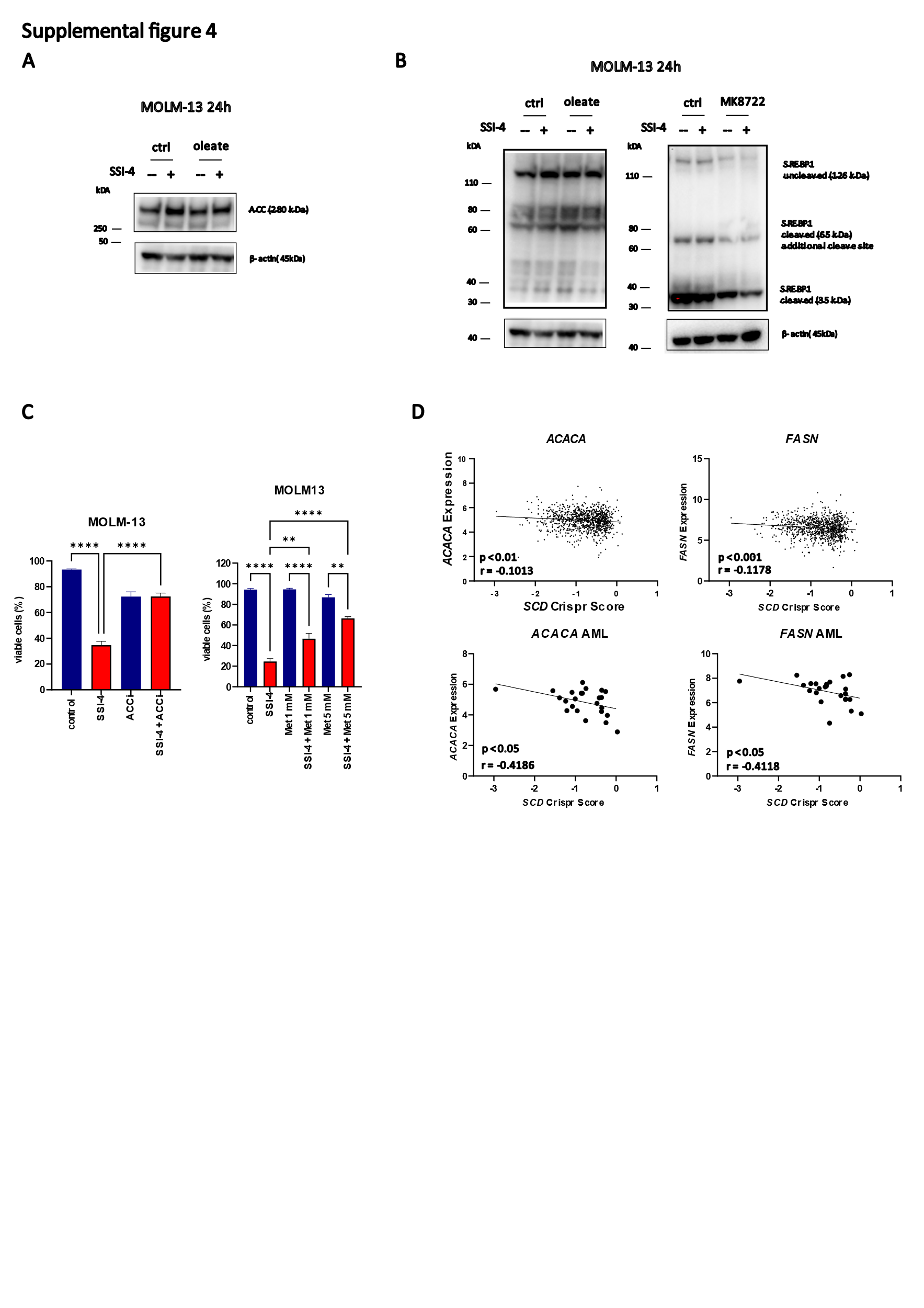

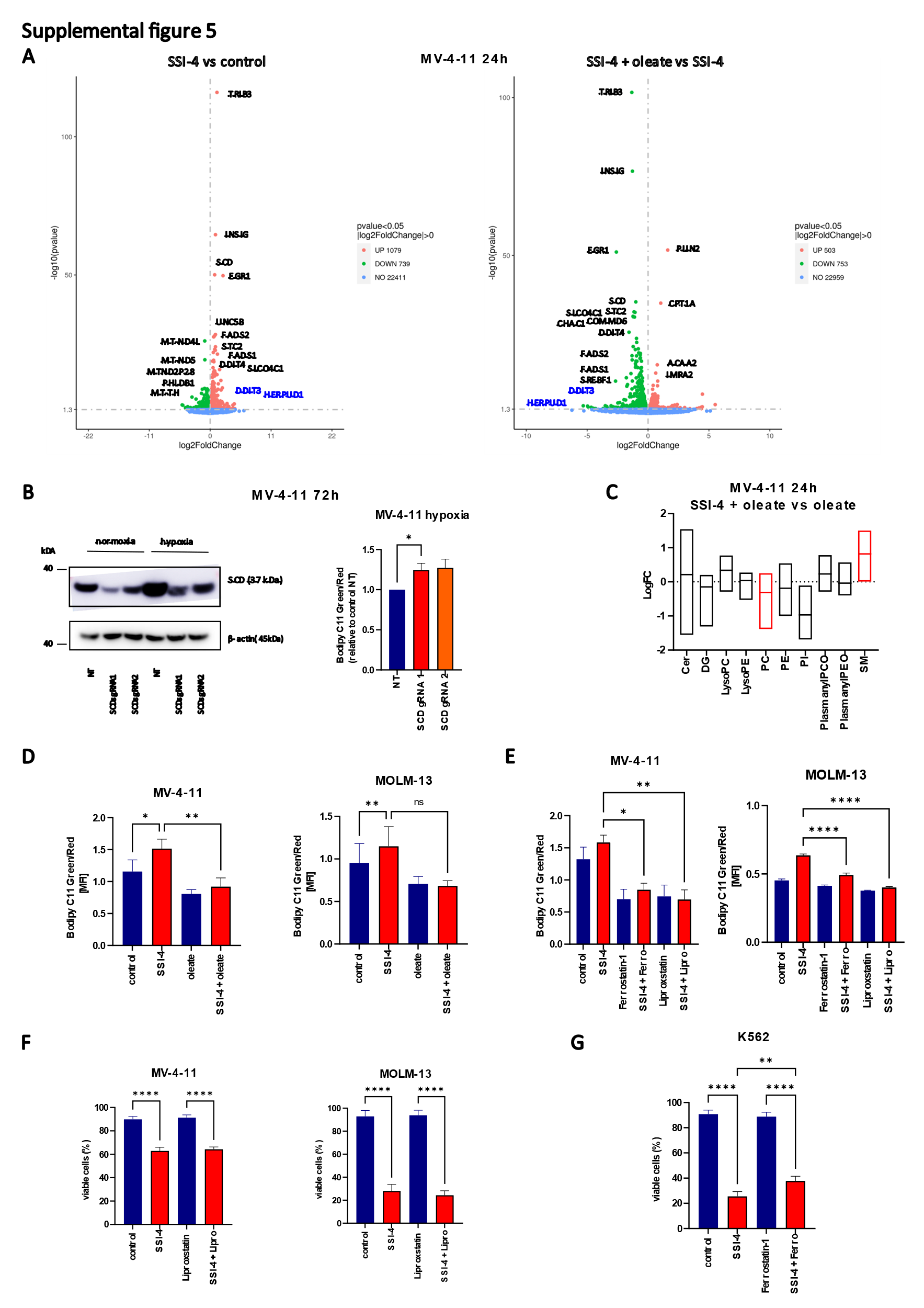

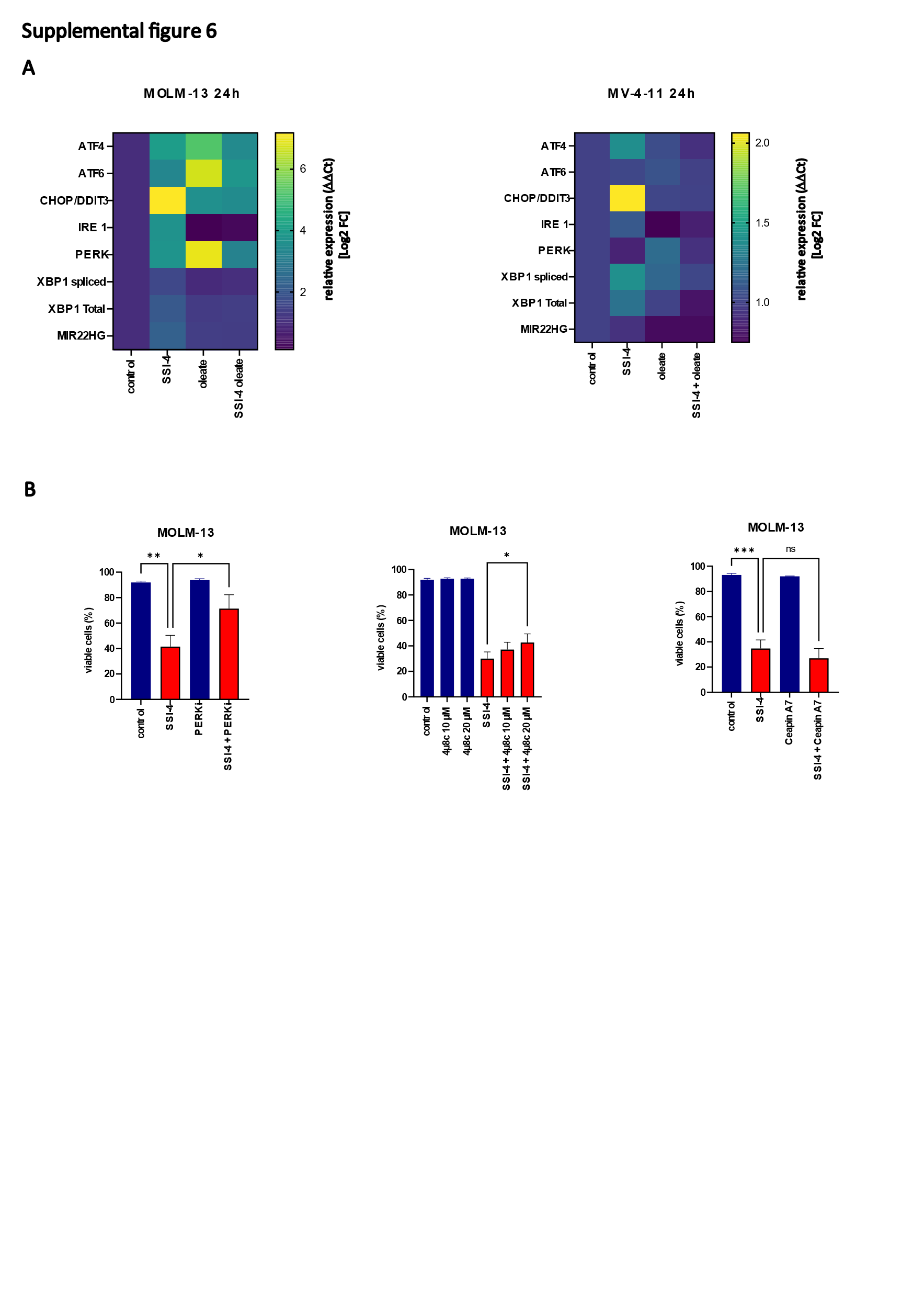

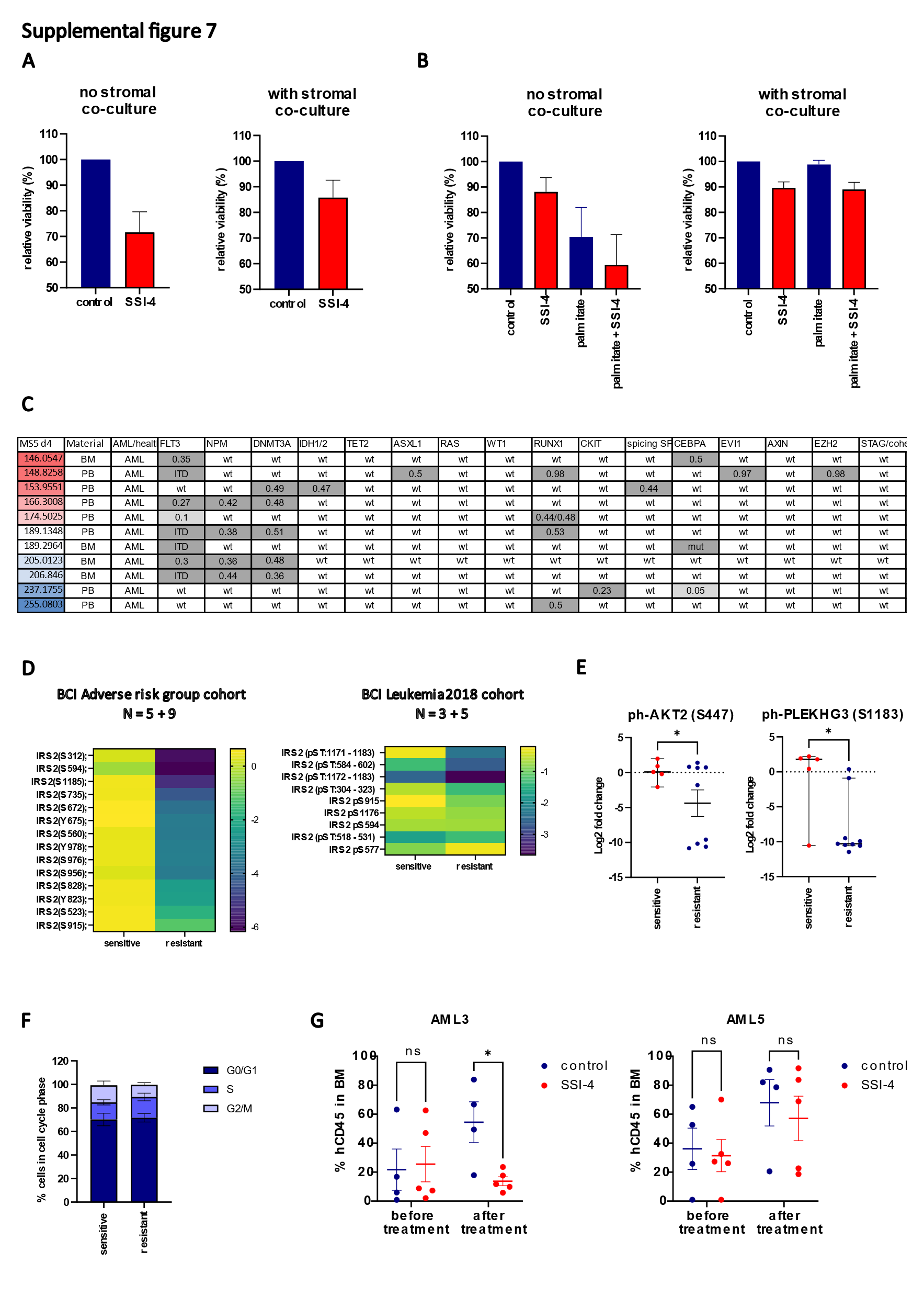

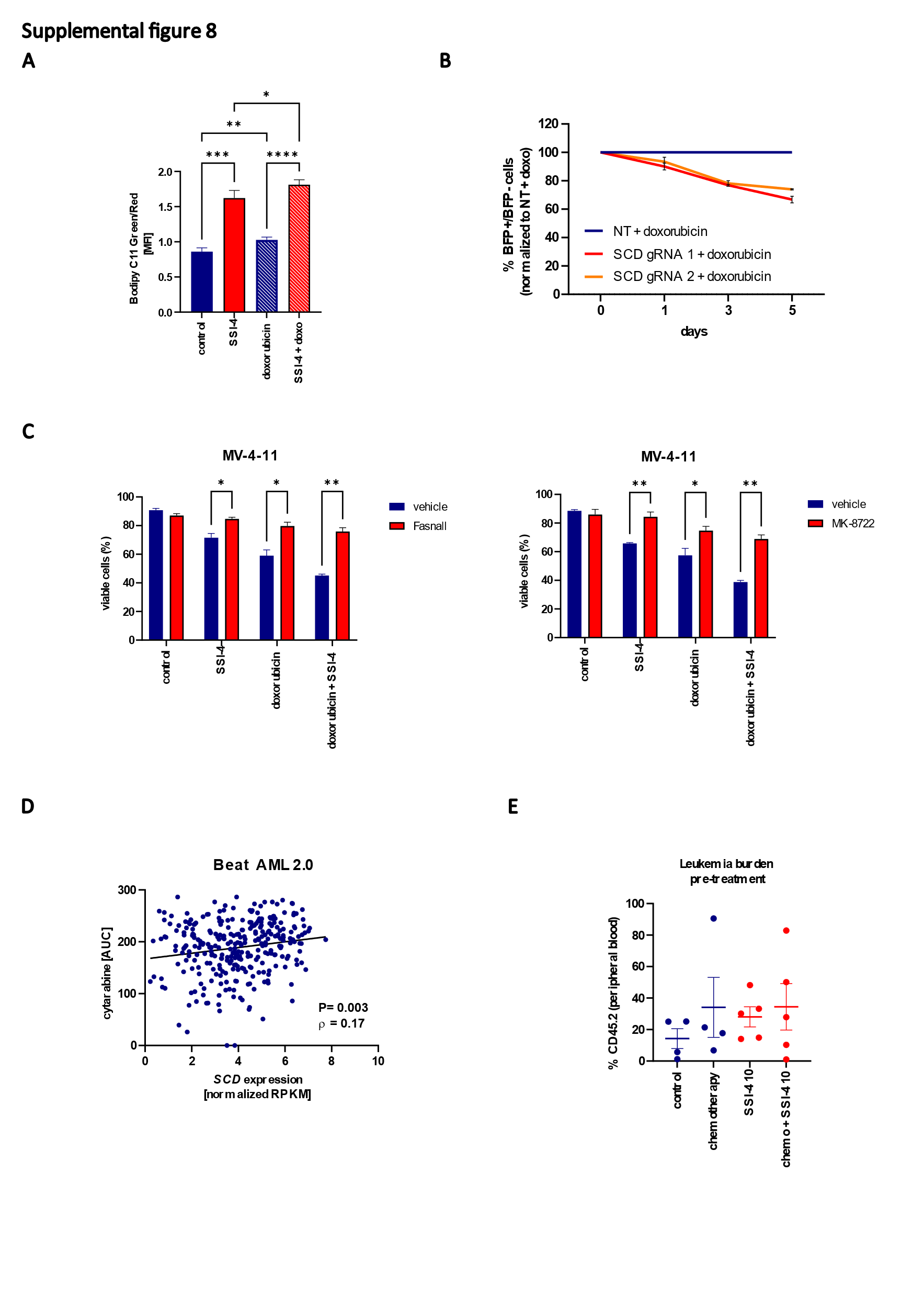

**Supplemental table 1. Reagents used in the study**

| **REAGENT** | **MANUFACTURER** | **CATALOGUE NUMBER** |
| --- | --- | --- |
| **Antibodies and dilutions (western blot)** | | |
| SCD Mouse mAb (CD.E10) (1:1000) | ThermoFisher | # MA5-27542, RRID: AB_2723611 |
| FASN Rabbit mAb (C20G5) (1:000) | Cell Signaling Technology | # 3180, RRID: AB_2100796 |
| ACC Rabbit mAb (C83B10) (1:1000) | Cell Signaling Technology | # 3676, RRID:AB_2219397 |
| AMPKα Rabbit mAb (D5A5) (1:1000) | Cell Signaling Technology | # 5831, RRID:AB_10622186 |
| ph-AMPKα (Thr172) Rabbit mAb (40H9) (1:1000) | Cell Signaling Technology | # 2535, RRID: AB_331250 |
| Cleaved PARP (Asp214) Rabbit mAb (D64E10) (1:1000) | Cell Signaling Technology | # 5625, RRID:AB_10699459 |
| Caspase-3 Rabbit mAb (D3R6Y) (1:1000) | Cell Signaling Technology | # 14220, RRID:AB_2798429 |
| PERK Rabbit mAb (C33E10) (1:1000) | Cell Signaling Technology | # 3192, RRID:AB_2095847 |
| eIF2α Rabbit mAb (D7D3) (1:1000) | Cell Signaling Technology | # 5324, RRID:AB_10692650 |
| Phospho-eIF2α (Ser51) Rabbit mAb (D9G8) (1:1000) | Cell Signaling Technology | # 3398, RRID:AB_2096481 |
| β-Actin Mouse mAb (8H10D10) | Cell Signaling Technology | # 3700, RRID:AB_2242334 |
| Anti-mouse IgG, HRP-linked Antibody | Cell Signaling Technology | # 7076, RRID:AB_330924 |
| Anti-rabbit IgG, HRP-linked Antibody | Cell Signaling Technology | # 7074, RRID:AB_2099233 |
| SREBP-2 Mouse Ab IgG-1C6 | BD Pharmingen™ | # 557037 RRID:AB_396560 |
| SREBP-1 Mouse mAb (2A4) | AbCam | # ab3259, RRID:AB_303650 |
| FADS2 Rabbit polyclonal Ab | AbCam | # ab232898 |
| **Antibodies and dilutions (flow cytometry)** |  |  |
| Annexin V FITC (1:30) | BioLegend | # 640945 |
| anti-human CD45 Pacific Blue™ (1:50) | BioLegend | # 982306, RRID:AB_2650649 |
| anti-mouse CD45 APC (1:50) | BioLegend | # 157605, RRID:AB_2876537 |
| anti-human CD19 Brilliant Violet 711™ (1:50) | BioLegend | # 302245, RRID:AB_2562062 |
| anti-human CD33 PE (1:50) | BioLegend | # 303404, RRID:AB_314348 |
| anti-mouse CD45.1 Brilliant Violet 711™ (1:50) | BioLegend | # 110739, RRID:AB_2562605 |
| anti-mouse CD45.2 FITC (1:50) | BioLegend | # 109805, RRID:AB_313442 |
| anti-mouse CD117 (c-kit) PE (1:50) | BioLegend | # 135105, RRID:AB_1877216 |
| anti-mouse Ly-6G/Ly-6C (Gr-1) PE/Cyanine7 (1:50, 1:1000) | BioLegend | # 108415, RRID:AB_313380 |
| anti-mouse/human CD11b APC (1:50, 1:1000) | BioLegend | # 101212, RRID:AB_312795 |
| anti-mouse/human CD45R/B220 PerCP (1:100) | BioLegend | # 103233, RRID:AB_893355 |
| anti-mouse CD19 APC/Cyanine7 | BioLegend | # 152411, RRID:AB_2922473 |
| anti-mouse CD8a PE (1:1000) | BioLegend | # 100707, RRID:AB_312746 |
| anti-mouse CD4 PE (1:5000) | BioLegend | # 100407, RRID:AB_312692 |
| Streptavidin Brilliant Violet 421™ (1:200) | BioLegend | # 405225 |
| anti-mouse Lineage Panel biotin (1:5) | BioLegend | # 133307, RRID:AB_11124348 |
| anti-mouse CD117 (c-Kit) Brilliant Violet 711™ (1:100) | BioLegend | # 105835, RRID:AB_2565956 |
| anti-mouse Ly-6A/E (Sca-1) APC/Cyanine7 (1:100) | BioLegend | # 108125, RRID:AB_10639725 |
| anti-mouse CD48 PE (1:250) | BioLegend | # 103405, RRID:AB_313020 |
| anti-mouse CD150 (SLAM) PE/Cyanine7 (1:100) | BioLegend | # 115914, RRID:AB_439797 |
| anti-human CD36 APC (1:20) | BD Pharmingen™ | # 550956, RRID:  AB_398480 |
| anti-human LDLR PE (1:20) | AbCam | # ab275614 |
| Human TruStain FcX™ (Fc Receptor Blocking Solution) | BioLegend | # 422302 RRID:AB_2818986 |
| TruStain FcX™ PLUS (anti-mouse CD16/32) Antibody | BioLegend | # 156604 |
| **Cytokines** | | |
| Recombinant Mouse IL-3 (carrier-free) | BioLegend | # 575504 |
| Recombinant Mouse IL-6 (carrier-free) | BioLegend | # 575704 |
| Recombinant Mouse SCF (carrier-free) | BioLegend | # 579704 |
| Recombinant Mouse G-CSF (carrier-free) | BioLegend | # 574604 |
| Recombinant Human G-CSF (carrier-free) | BioLegend | # 578604 |
| Recombinant Human IL-3 (carrier-free) | BioLegend | # 578004 |
| Recombinant Human TPO (carrier-free) | BioLegend | # 763704 |
| **Bacterial and virus strains** | | |
| E.coli DH5α | Kind gift of B. Huntly | NCBI:txid668369 |
| psPAX2 | Addgene | #12260, RRID:Addgene_12260 |
| pMD2.G | Addgene | #12259, RRID:Addgene_12259 |
| MSCV-Hoxa9-neo | Kind gift of T. Sommerville | NA |
| MSCV-Meis1a-puro | Kind gift of T. Sommerville | NA |
| **Chemicals** | | |
| Propidium iodide solution | Sigma-Aldrich | # P4864 |
| TO-PRO-3 | Life Technologies | # T3605 |
| 7-AAD | BioLegend | # 420404 |
| Annexin Binding Buffer | bioWORLD | # 21720002 |
| Ammonium Chloride Solution | Stemcell Technologies | # 07850 |
| SSI-4 | Modulation Therapeutics Inc. | NA |
| A939572 | Sigma-Aldrich | # SML2356 |
| Sodium palmitate | Sigma-Aldrich | # P9767 |
| Sodium oleate | Sigma-Aldrich | # O7501 |
| Fasnall (benzenesulfonate) | Cayman Chemical | # 19957 |
| PF-05175157 | MedChemExpress | # HY-12942 |
| MK-8722 | MedChemExpress | # HY-111363 |
| Ferrostatin-1 | MedChemExpress | # HY-100579 |
| Liproxstatin-1 | MedChemExpress | # HY-12726 |
| Q-VD-OPh | MedChemExpress | # HY-12305 |
| GSK2656157 | Sigma-Aldrich | # 504651 |
| 4μ8C | Sigma-Aldrich | # SML0949 |
| Doxorubicin hydrochloride | Sigma-Aldrich | # D1515 |
| 1-β-D-Arabinofuranosylcytosine (cytarabine) | Sigma-Aldrich | # 251010 |
| Puromycin dihydrochloride | Sigma-Aldrich | # P8833 |
| Doxycycline hyclate | Sigma-Aldrich | # D5207 |
| Sucrose | Sigma-Aldrich | # 573113 |
| D-GLUCOSE (U-13C6, 99%) | Cambridge Isotope Laboratories Inc. | # CLM-1396-PK |
| NuPAGE™ 4 to 12%, Bis-Tris, 1.0–1.5 mm, Mini Protein Gels | Invitrogen™ | # NP0321BOX |
| NuPAGE™ 3 to 8%, Tris-Acetate, 1.0–1.5 mm, Mini Protein Gels | Invitrogen™ | # EA0375BOX |
| NuPAGE™ LDS Sample Buffer (4X) | Invitrogen™ | # NP0007 |
| NuPAGE™ MES SDS Running Buffer (20X) | Invitrogen™ | # NP0002 |
| NuPAGE™ Tris-Acetate SDS Running Buffer (20X) | Invitrogen™ | # LA0041 |
| NuPAGE™ Transfer Buffer (20X) | Invitrogen™ | # NP00061 |
| Immobilon®-P PVDF Membrane | Millipore | # IPVH00005 |
| Novex™ Sharp Pre-stained Protein Standard | Invitrogen™ | # LC5800 |
| PageRuler™ Plus Prestained Protein Ladder, 10 to 250 kDa | ThermoFisher™ | # 26619 |
| Protease Inhibitor Cocktail Set I | Sigma-Aldrich | # 539131 |
| Phosphatase Inhibitor Cocktail Set III | Millipore | # 524627 |
| Clarity Western ECL Substrate | BioRad | # 1705061 |
| SuperSignal™ West Pico PLUS Chemiluminescent Substrate | ThermoFisher™ | # 34579 |
| EasySep™ Human TCR Alpha/Beta Depletion Kit | StemCell Technologies | # 17847 |
| TransIT®-Lenti Transfection Reagent | Mirus Bio | # MIR 6603 |
| Polybrene Infection / Transfection Reagent | Sigma-Aldrich | # TR-1003 |
| **Assays** | | |
| Zombie NIR™ Fixable Viability Kit | BioLegend | # 423106 |
| BODIPY™ 581/591 C11 (Lipid Peroxidation Sensor) | Invitrogen™ | # D3861 |
| C1- BODIPY™ 500/510 C12 (Lipid Uptake Sensor) | Invitrogen™ | # D3823 |
| Direct-zol RNA Microprep | Zymo Research | # R2061 |
| High-Capacity cDNA Reverse Transcription Kit | Applied Biosystems™ | # 4368814 |
| PowerUp™ SYBR™ Green Master Mix | Applied Biosystems™ | # A25741 |
| EndoFree Plasmid Maxi Kit (10) | Qiagen | # 12362 |
| **Oligonucleotides** | | |
| SCD gRNA 1: 5’-CACCGATATATGACCCCACCTACA-3’ | Sigma-Aldrich | NA |
| SCD gRNA 2: 5’-CACCGCATATTCAACCTTGGGGCT-3’ | Sigma-Aldrich | NA |
| NT gRNA: 5’-CACCGATTTTCGTACCCTGGGACGC-3’ | Sigma-Aldrich | NA |
| pKLV2 primer: 5’-AGATAATTAGAATTAATTTGACTG-3’ | Sigma-Aldrich | NA |
| **Recombinant DNA** | | |
| lentiCas9-Blast | Addgene | #52962, RRID:Addgene_52962 |
| pKLV2-U6gRNA5(BbsI)-PGKpuro2ABFP-W | Addgene | #67974, RRID:Addgene_67974 |
| **Media** | | |
| MyeloCult™ H5100 | StemCell Technologies | # 05150 |
| MethoCult™ GF M3434 | StemCell Technologies | # 03434 |
| MethoCult™ M3231 | StemCell Technologies | # 03231 |
| RPMI | Gibco™ | # 11875093 |
| DMEM | Gibco™ | # 11965092 |
| IMDM | Gibco™ | # 12440053 |
| MEM-α | Gibco™ | # 12571063 |
| OptiMEM | Gibco™ | # 31985070 |

**Supplemental table 2. Primers for human cDNA (qPCR) used in the study**

|  | **Forward** | **Reverse** |
| --- | --- | --- |
| **ATF4** | GCTAAGGCGGGCTCCTCCGA | ACCCAACAGGGCATCCAAGTCG |
| **ATF6** | ATGAAGTTGTGTCAGAGAACC | CTCTTTAGCAGAAAATCCTAG |
| **CHOP** | GGAGCATCAGTCCCCCACTT | TGTGGGATTGAGGGTCACATC |
| **IRE 1** | AGTCAGTTCTGCGTCCGCT | TGGTACTTCCAAAAATCCCGAGG |
| **PERK** | ATGCTTTCACGGTCTTGGTC | TCATCCAGCCTTAGCAAACC |
| **XBP1 spliced** | TTGCTGAAGAGGAGGCGGAA | CTGCACCTGCTGCGGACTCAG |
| **XBP1 total** | TTCCGGAGCTGGGTATCTCA | GAAAGGGAACCCCCGTATCC |
| **MIR22HG** | CCTCGTGCAGCAACCCC | GTGAGGGCGTGAGAGGAAC |
| **actB** | GCCGCCAGCTCACCAT | TCGTCGCCCACATAGGAATC |

**Supplemental data 1. Lipidomics analysis raw data**

**Supplemental data 2. Fatty acids profiling cell lines raw data**

**Supplemental data 3. Fatty acids profiling primary samples raw data**
